# Cell state diversity promotes metastasis through heterotypic cluster formation in melanoma

**DOI:** 10.1101/2020.08.24.265140

**Authors:** Nathaniel R. Campbell, Anjali Rao, Maomao Zhang, Maayan Baron, Silja Heilmann, Maxime Deforet, Colin Kenny, Lorenza Ferretti, Ting-Hsiang Huang, Manik Garg, Jérémie Nsengimana, Emily Montal, Mohita Tagore, Miranda Hunter, Julia Newton-Bishop, Mark R. Middleton, Pippa Corrie, David J. Adams, Roy Rabbie, Mitchell P. Levesque, Robert A. Cornell, Itai Yanai, Joao B. Xavier, Richard M. White

**Author notes:** Novo Nordisk Foundation Center for Stem Cell Biology, University of Copenhagen, 2200 Copenhagen N, Denmark (Silja Heilmann). Sorbonne Université, CNRS, Institut de Biologie Paris-Seine (IBPS), Laboratoire Jean Perrin (LJP), F-75005, Paris, France. Co-communicating co-senior authors: J.B.X. and R.M.W.

## Abstract

In melanoma, transcriptional profiling has revealed multiple co-existing cell states, including proliferative versus invasive sub-populations that have been posited to represent a “go or grow” tradeoff. Both of these populations are maintained in tumors, but how they physically interact to promote metastasis is unknown. We demonstrate that these subpopulations form spatially structured heterotypic clusters that cooperate in the seeding of metastasis. We unexpectedly found that INV cells were tightly adherent to each other, and formed clusters with a rim of PRO cells. Intravital imaging demonstrated cooperation between these populations, in which the INV cells facilitated the spread of less metastatic PRO cells. We identified the TFAP2 neural crest transcription factor as a master regulator of both clustering and the PRO/INV states. Our data suggest a framework for the co-existence of these two divergent cell populations, in which differing cell states form heterotypic clusters that promote metastasis via cell-cell cooperation.

## INTRODUCTION

### Cell state heterogeneity in cancer

Tumors are heterogenous populations of cells that contain a variety of subpopulations differing both through genetic and non-genetic mechanisms (Hinohara and Polyak, 2019). One such form of heterogeneity is transcriptional. Numerous studies using bulk or single-cell transcriptomics have demonstrated the existence of transcriptional subpopulations of cells, often referred to as cancer cell states (Hinohara and Polyak, 2019; Hoek and Goding, 2010). The mechanisms that generate the different cell states, and how those states interact with each other remains poorly understood.

### Melanomas have multiple co-existing cell states, including PRO/INV cells

Melanoma has long been noted to exhibit a wide range of phenotypic properties such as pigmentation and invasiveness (Houghton et al., 1987), which is related to underlying transcriptional heterogeneity. Such heterogeneity was initially studied by bulk RNA microarray (Bittner et al., 2000; Hoek et al., 2006; Widmer et al., 2012) and RNA-sequencing technologies (Rambow et al., 2015; Verfaillie et al., 2015), but more recent single-cell RNA sequencing (Tirosh et al., 2016; Wouters et al., 2020) has increased the granularity of these distinctions. Increasing evidence points to at least four distinct cell states (Rambow et al., 2018; Tsoi et al., 2018; Wouters et al., 2020), with the most consistently identified ones comprising a proliferative (PRO) versus invasive (INV) cell state. Individual cells tend to be PRO or INV, but not both (Hoek et al., 2008; Hoek et al., 2006; Rambow et al., 2015; Tirosh et al., 2016; Verfaillie et al., 2015; Widmer et al., 2012)—a tradeoff reminiscent of the “grow or go” hypothesis (Hatzikirou et al., 2010; Matus et al., 2015). The PRO vs. INV populations have been tightly linked to the process of phenotype switching, a phenomenon in which cells can bidirectionally move between these two PRO vs. INV extremes after induction by signals such as Wnt5A, EDN3, hypoxia, inflammation or nutrient deprivation from the microenvironment (Carreira et al., 2006; Eichhoff et al., 2010; Falletta et al., 2017; Hoek et al., 2008; Hoek et al., 2006; Kim et al., 2017; Pinner et al., 2009; Weeraratna et al., 2002; Widmer et al., 2012). The PRO vs. INV state is in part controlled by the melanocyte master transcription factor MITF (Carreira et al., 2006; Eichhoff et al., 2010), with the PRO cells generally being MITF^HI^ and INV cells being MITF^LO^, although many other genes such as AXL have been linked to these states (Tirosh et al., 2016; Verfaillie et al., 2015). Some data posits the existence of biphenotypic cells, with individual cells having characteristics of both PRO and INV cells upon deletion of *Smad7* (Tuncer et al., 2019). The extent to which these states phenotype switch, or remain relatively fixed in their identities, has important implications in whether we should be targeting the plasticity itself or the states themselves.

### Functions of co-existing cell states in tumor evolution

Despite evidence that these and other (Baron et al., 2020; Rambow et al., 2019; Rambow et al., 2018; Tsoi et al., 2018) subpopulations exist in tumors, little is known about how these states co-exist within the tumor, or whether they cooperate to promote tumorigenic phenotypes such as metastasis. While some cell states (i.e. a neural crest-like population driven by RXRG or via sensitivity to iron-dependent ferroptotic cell death) have been clearly linked to resistance to MAPK inhibitor therapy (Rambow et al., 2018; Tsoi et al., 2018), the role of the PRO/INV populations has been best studied in the context of metastasis. Analogous to an EMT-like process in epithelial cancers, it was hypothesized that PRO cells could phenotype switch to a more INV state by molecules such as Wnt5A (Weeraratna et al., 2002), and become more migratory and metastatic. While this switching model likely explains metastases in some patients, it does not fully explain why these cell state subpopulations seem to co-exist, albeit at different ratios, in nearly all patients examined. Mixing PRO with INV cells (albeit from different patients) was shown to lead to polyclonal metastatic seeding (Chapman et al., 2014; Rowling et al., 2020), raising the possibility that different cell states, each with distinct phenotypes, might cooperate with each other to promote phenotypes such as metastasis.

### Cooperation between cell states as a mechanism for metastasis

Cooperation—a social behavior where one individual increases the fitness of another—is widely studied in the contexts of ecology and evolution (Ågren et al., 2019; Archetti and Pienta, 2019; Foster, 2011; Hauser et al., 2009; Korolev et al., 2014). Its potential role in cancer remains understudied, especially *in vivo*. In a Wnt1-driven mouse model of breast cancer both basal and luminal cell types emerge during tumorigenesis; both populations are required for tumor growth, with Wnt1 produced by the luminal cells supporting growth of the basal population (Cleary et al., 2014). This model demonstrated how cooperation can provide a selective pressure for the maintenance of heterogeneity within tumors. Along the same lines, experiments with heterotypic tumors where subpopulations overexpressed factors previously implicated in tumor progression revealed a minor subclone capable of driving enhanced proliferation of the entire tumor (Marusyk et al., 2014). This clone acted by secreting IL-11 to stimulate vascular growth and reorganization of the extracellular matrix. When this clone was combined with a clone expressing FIGF, the otherwise nonmetastatic tumors gained the ability to metastasize. How these different cell states physically interact, and the mechanism by which they might cooperate in metastasis, remain unknown.

Here, show that PRO and INV cell states can form heterotypic clusters that cooperate in metastasis. Circulating tumor cell clusters have long been recognized as a particularly potent mechanism for metastasis (Aceto et al., 2014; Fidler, 1973; Glaves, 1983; Liotta et al., 1976; Long et al., 2016; Luo et al., 2014; Mayhew and Glaves, 1984; Watanabe, 1954), and are strongly associated with worse outcome. Using a transgenic zebrafish model of melanoma (Ceol et al., 2011; Patton et al., 2005; White et al., 2011), we show that PRO and INV transcriptional states spontaneously aggregate into spatially ordered clusters, with a rim of PRO cells surrounding a dense core of INV cells. Unexpectedly, we find that the more INV cells express higher levels of adhesion molecules, a finding recapitulated in human melanoma specimens. These heterotypic clusters recapitulate developmental adhesive sorting, in which embryonic cells with differential levels of adhesion proteins spontaneously form similar structures. Consistent with this notion, we find that this cluster structure is regulated by the developmental neural crest transcription factor TFAP2, which mediates the PRO vs. INV state and metastatic seeding capacity. While phenotype switching is a likely mechanism of metastasis in some patients, our data provide an alternative mechanism by which relatively fixed cell states physically cooperate to promote metastasis via cooperative clustering of divergent cell states.

## RESULTS

### Characterization of PRO/INV cell states

To address this question, we utilized a zebrafish model of melanoma that allows for longitudinal single cell analysis of these heterogeneous subpopulations in metastasis formation (Cagan et al., 2019; Heilmann et al., 2015). From a transgenic melanoma in a BRAF^V600E^;p53^-/-^ animal (Ceol et al., 2011; Kaufman et al., 2016; Patton et al., 2005; White et al., 2011) we generated a low-passage zebrafish melanoma cell line, ZMEL1 (Heilmann et al., 2015), and phenotyped multiple cultures to identify populations enriched for either proliferative (ZMEL1-PRO) or invasive (ZMEL1-INV) phenotypes (Figure 1a). Consistent with the previous characterization of PRO and INV states (Widmer et al., 2012), we observed a small but consistent proliferation difference, and a more substantial motility difference, between the ZMEL1-PRO and -INV states (Figure 1b-c, Figure S1a-b). To confirm that this recapitulates human PRO and INV states, we performed RNA-sequencing analysis (RNA-seq) on these two ZMEL1 populations and found a strong association between ZMEL1-INV and -PRO states and published human INV and PRO gene signatures (Hoek et al., 2006; Tirosh et al., 2016; Verfaillie et al., 2015; Widmer et al., 2012), respectively, with the INV signature from Hoek et al. (Hoek et al., 2006) the top gene set (Figure 1d-e, Figure S1c, Supplementary Tables 1,6,7).

**Figure 1.**
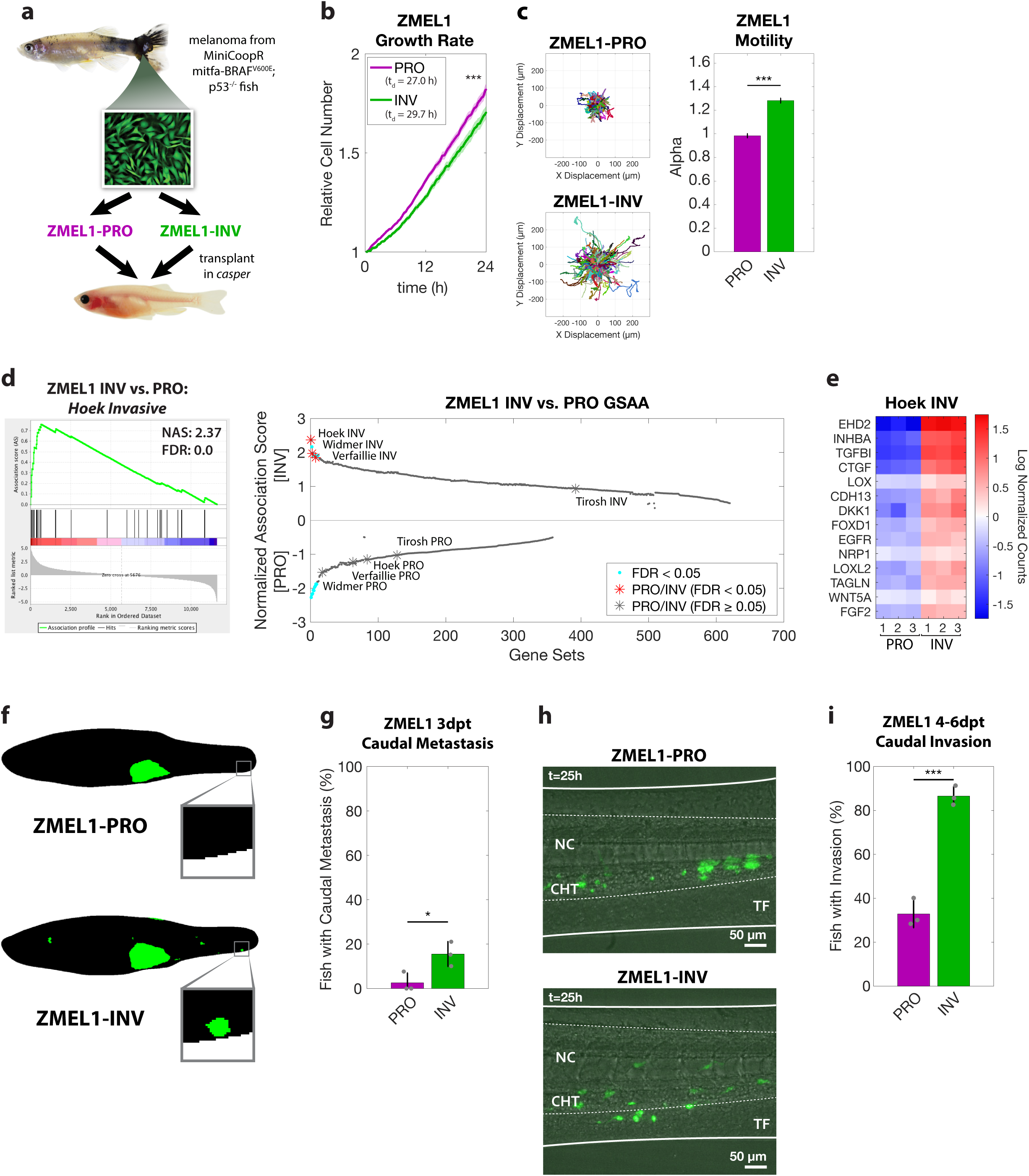
PRO and INV coexist in zebrafish melanoma, with INV cells metastasizing more frequently due to increased extravasation. **a**. Proliferative (PRO) and invasive (INV) subpopulations were identified from the ZMEL1 zebrafish melanoma cell line, which was originally isolated from a transgenic zebrafish and can be transplanted into transparent *casper* zebrafish (adapted with permission from (Heilmann et al., 2015)). **b-c**. Tracking of individual cells by time-lapse microscopy (both p<0.001 by linear regression, N=4 independent experiments). **b**. Growth curves (mean ± SE of mean, smoothed with moving window average of 5 time points) and doubling time (mean [95% CI]: 27.0 h [26.9, 27.1] vs. 29.7 h [29.6, 29.9] for ZMEL1-PRO and -INV, respectively. **c**. (left) Representative displacements of 500 tracks, and (right) model estimates ± 95% CI for alpha, the slope of the log-log plot of mean squared displacement vs. lag time (tau) for each ZMEL1-PRO and -INV. Larger alpha indicates more persistent motion. **d**. (left) The INV signature from Hoek et al. (Hoek et al., 2006) was the top gene set by Gene Set Association Analysis (GSAA) of ZMEL1-INV vs. -PRO RNA-seq. (Right) Dual waterfall plot of GSAA ranked by false discovery rate (FDR). Literature PRO/INV gene sets are indicated with an asterisk and colored according to FDR. **e**. Heatmap of genes in Hoek INV signature that are differentially expressed between ZMEL1-PRO and -INV (log_2_ fold change cutoff ± 1.5, p_adj_ < 0.05). Human ortholog gene names are used for clarity (see Figure S1e for zebrafish gene names). **f**. Segmentation of representative images of ZMEL1-PRO and -INV tumors and distant metastases (e.g. to caudal region [box]) at 3 days post-transplant (3dpt). Original images shown in Figure S1e. **g**. Quantification of caudal metastases seeded by ZMEL1 populations at 3 dpt (OR [95% CI]: 11.52 [1.41, 93.73]; p=0.022 by logistic regression; N=3 independent experiments with PRO/INV 9/10, 31/33, and 13/13 fish per group, respectively; n=109 fish total; plot shows mean ± SD). **h**. Representative images at 1 dpt from time lapse confocal microscopy of ZMEL1 cells transplanted intravenously in larval zebrafish. Arrowhead indicates group of cells invading from the notochord (NC) and caudal hematopoietic tissue (CHT) into the tail fin mesenchyme (TF). Images are representative of n=13 fish per cell type. See Supplementary Video 1 for full time lapse. **i**. Quantification of caudal tissue invasion by imaging at 4-6dpt (N=3 independent experiments with PRO/INV 23/23, 21/21, and 19/23 fish per group, respectively; OR [95% CI]: 13.58 [5.56, 33.18]; p<0.001 by logistic regression, plot shows mean ± SD).

To compare the metastatic potential of ZMEL1-PRO and ZMEL1-INV, we transplanted each population orthotopically into the subcutaneous tissue of transparent *casper* zebrafish and followed their growth and metastasis by whole-fish fluorescence microscopy (Heilmann et al., 2015) (Figure 1f-g, Figure S1d-e). Fish harboring ZMEL1-INV tumors were significantly more likely to have distant metastases three days post-transplant (3 dpt), particularly in the caudal region of the fish, an anatomical region relatively resistant to metastasis (Heilmann et al., 2015). To investigate this difference in detail, we transplanted each population intravenously in larval *casper* zebrafish where we followed the seeding of metastases by confocal time-lapse microscopy. ZMEL1-INV cells extravasated more effectively than ZMEL1-PRO cells within the first dpt (Figure 1h, Supplementary Video 1). To quantify this difference, we tracked metastatic progression in similarly transplanted larval fish by daily whole-fish imaging; ZMEL1-INV cells invaded into the caudal tissue in a significantly higher proportion of fish at the experiment endpoint (4-6 dpt, Figure 1i). Since the cells were injected intravenously, these findings implicate extravasation as a key step of metastatic spread where INV cells are more effective than PRO cells.

### PRO/INV cells form heterotypic clusters

To identify functional processes differentiating PRO and INV populations, we performed Gene Ontology (GO) association analysis on our RNA-seq data. This analysis unexpectedly revealed a strong association between the INV state and signatures of enhanced cell-cell adhesion, with many adhesion genes upregulated (Figure 2a-b, Figure S2a-c, Supplementary Tables 6,7). This association was surprising, as the classical model of cell invasion involves the loss of cell adhesion and the gain of individual motility, the opposite of what we observed (Gupta et al., 2005; Li et al., 2015; Padmanaban et al., 2019). To test this paradoxical finding, we utilized a three-dimensional (3D) cluster formation assay in low-attachment plates. Under these conditions, while the ZMEL1-PRO cells tended to stay as individual cells or small clusters, the ZMEL1-INV population formed strikingly large, spherical clusters (Figure 2c-d Supplementary Video 2) in agreement with increased adhesive properties. To test whether the association between invasiveness and cell clustering is a general feature of melanoma, we compared the INV signature defined by Hoek et al. (Hoek et al., 2006) with that of the cell-cell adhesion genes most associated with ZMEL1-INV in a panel of 56 melanoma cell lines available in the Cancer Cell Line Encyclopedia (CCLE) (Ghandi et al., 2019) and 472 clinical melanoma samples from The Cancer Genome Atlas (TCGA) (Cancer Genome Atlas Network, 2015). In both cohorts, the expression of cell-cell adhesion genes correlated strongly with the INV cell state (Figure S2d-e). To validate this finding functionally, we assayed the cluster-forming ability of a panel of nine human melanoma cell lines. We observed a strong correlation between cluster formation and the INV state, consistent with our zebrafish findings (Figure 2e). Taken together, these results indicate that melanomas that are invasive and metastatic tend to form cluster aggregates.

**Figure 2.**
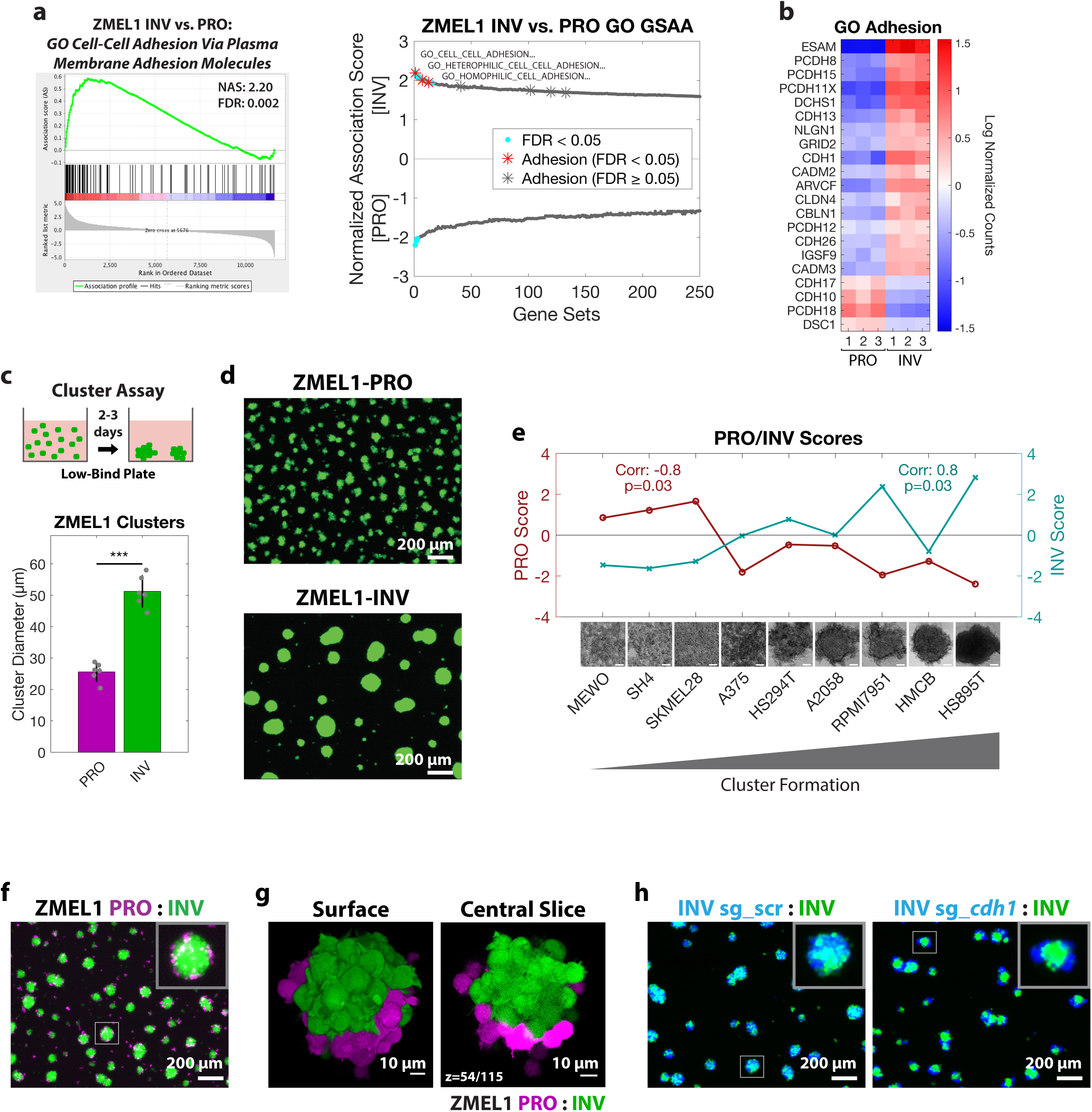
Cluster formation by INV state drives spatial patterning of melanoma clusters. **a**. (left) Plot of top Gene Ontology (GO) gene set by GSAA for ZMEL1-INV vs. -PRO RNA-seq. (right) Dual waterfall plot of top/bottom 250 gene sets from GO analysis (for full plot see Figure S2a). Adhesion GO gene sets are indicated with an asterisk and colored according to false discovery rate (FDR) **b**. Heatmap of genes in adhesion GO gene sets (FDR < 0.05, n=3) that are differentially expressed between ZMEL1-PRO and -INV (log_2_ fold change cutoff ± 1.5, p_adj_ < 0.05). Human ortholog gene names are used for clarity (see Figure S2b-c for absolute expression and zebrafish gene names). **c**. (top) Schematic of assay and (bottom) quantification of cluster formation in low-bind plates after 2 days (N=6 independent experiments, p<0.001 by two-sided t-test, plot shows mean ± SD). **d**. Representative images of clusters formed after 3 days. **e**. Human melanoma cell lines ranked by cluster forming ability (left to right: low to high) with PRO/INV gene expression scores from Hoek et al. (Hoek et al., 2006) (Spearman correlation and Bonferroni-corrected p-values shown, scale bar 100μm). **f**. Co-clusters of 1:1 mixture of ZMEL1-PRO and - INV. **g**. (left) 3D opacity rendering and (right) central slice (slice 54 of 115) of confocal imaging through co-cluster of ZMEL1-PRO and ZMEL1-INV. **h**. Co-clusters of 1:1 mixture of ZMEL1-INV and ZMEL1-INV with either control (sg_scr) or *cdh1* (sg_*cdh1*) sgRNA.

Individual primary patient melanomas comprise both PRO and INV subpopulations, and disseminated metastases preserve that diversity (Tirosh et al., 2016), raising the question of whether these subpopulations interact. Circulating tumor cell (CTC) clusters—comprised either of tumor cells or tumor and microenvironmental cells—are increasingly recognized for their role in promoting metastatic spread, facilitating diversity at metastatic sites (Aceto et al., 2014; Cheung et al., 2016; Gkountela et al., 2019; Maddipati and Stanger, 2015; Szczerba et al., 2019). Because the ZMEL1-PRO and -INV populations were isolated from a single primary tumor, we sought to establish whether the two could interact in clusters. Differential labeling of the PRO vs. INV cells revealed that the two cell states consistently generated co-clusters with a coherent spatial structure, with ZMEL1-INV cells at the core and ZMEL1-PRO cells at the rim, reminiscent of developmental cadherin sorting (Foty and Steinberg, 2005) (Figure 2f-g, Figure S2f-g, Supplementary Video 3). Indeed, CRISPR/Cas9 induced deletion of *cdh1* in ZMEL1-INV partially phenocopied ZMEL1-PRO, both decreasing the cluster size relative to INV clusters and causing spatial sorting of mixed clusters (Figure 2h, Figure S3a-c). Deletion of *cdh1* alone was insufficient, however, to induce changes in the metastatic rate of ZMEL1-INV (Figure S3d-e), suggesting that a broader set of adhesion and invasion genes, and not only *cdh1*, underlies the observed phenotypes. This stereotyped spatial organization of ZMEL1-PRO and -INV clusters motivated us to investigate whether interaction between these two populations would play a role *in vivo*.

### PRO/INV heterotypic clusters cooperate in metastasis

To assay these interactions *in vivo* during metastatic dissemination, we transplanted a 1:1 mixture of the ZMEL1-PRO and -INV populations intravenously in zebrafish larvae and followed them by confocal time-lapse microscopy (Figure 3a, Supplementary Video 4). We observed the spontaneous formation of intravascular tumor cell clusters comprised of cells from one or both cell states, consistent with both previous intravital imaging (Liu et al., 2018) and the detection of heterogenous CTC clusters in the blood of melanoma patients (Khoja et al., 2014). More notably, we observed that nearly half (11 out of 24) of ZMEL1-PRO extravasation events were co-extravasations of heterotypic tumor cell clusters (Figure S4a). We detected a pattern of collective motility suggesting that cells from the same heterotypic cluster extravasated collectively, with ZMEL1-INV cells behaving as leader cells and ZMEL1-PRO as followers. These detailed observations suggest that the PRO and INV states known to coexist in primary tumors can form heterotypic clusters and interact in the seeding of metastases.

**Figure 3.**
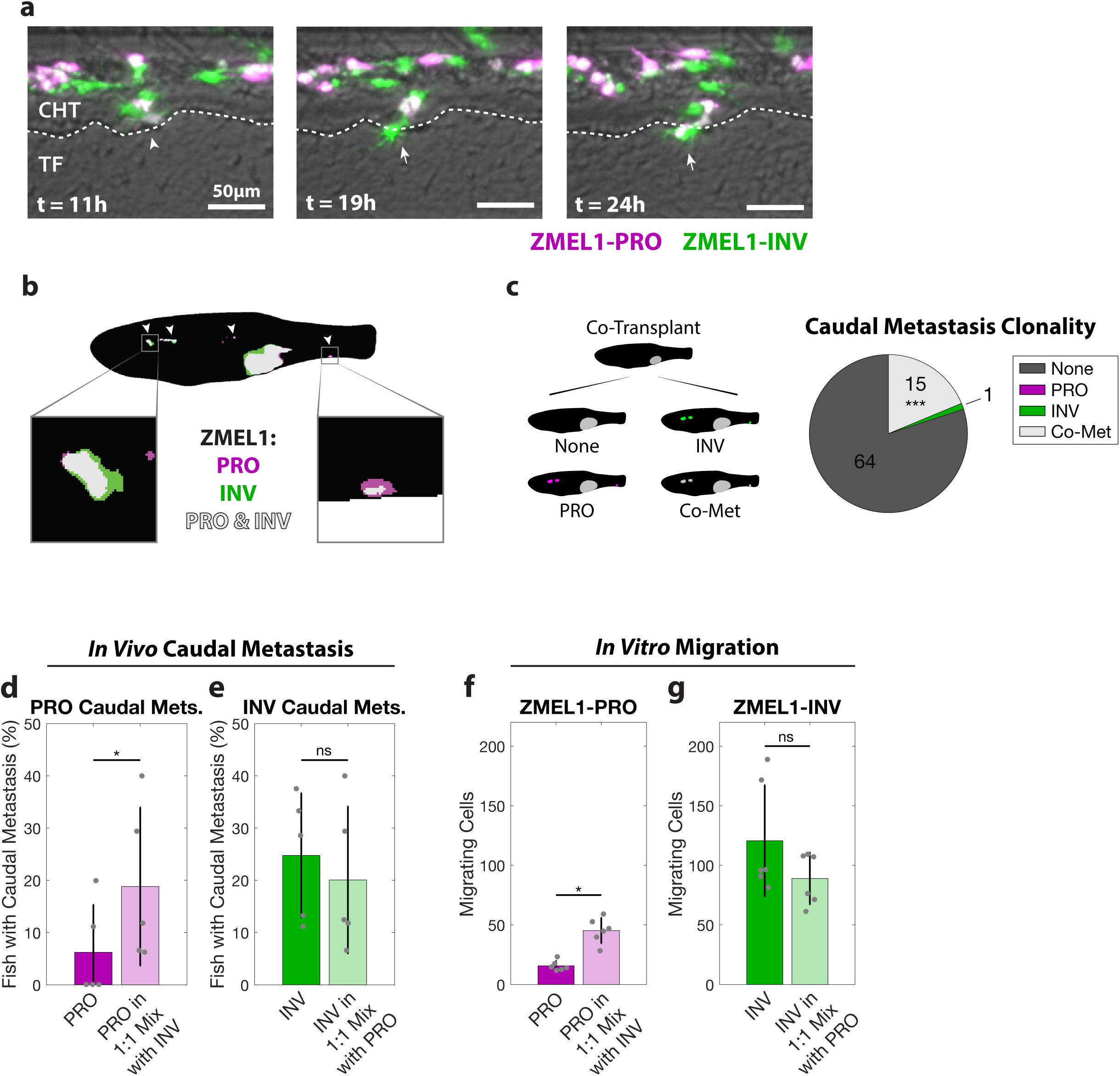
PRO and INV cooperate in metastasis via co-extravasation. **a**. In the first 24 hours following intravenous transplant of ZMEL1-PRO and -INV, a mixed cluster of both populations (left, arrowhead) extravasated from the caudal hematopoietic tissue (CHT) into the tail fin mesenchyme (TF)—with ZMEL1-INV leading (middle, arrow) and ZMEL1-PRO following (right, arrow). **b**. Segmented representative image of adult zebrafish with orthotopic transplant of 1:1 mixture of ZMEL1-PRO and -INV. Arrowheads indicate polyclonal metastases, including to the kidney and caudal regions (left and right boxes, respectively). Original image shown in Figure S4b. **c**. Number of fish co-transplanted with a 1:1 mixture of ZMEL1-PRO and - INV that have no caudal metastases (None), caudal metastases comprised of exclusively PRO or INV, or caudal metastases formed by co-metastasis (Co-Met) of both cell types (N=5 independent experiments with 17, 15, 16, 15, and 17 fish each; 80 fish total; p<0.001 by Mantel-Haenszel’s test for null hypothesis of no interaction). **d**. Percentage of fish with ZMEL1-PRO caudal metastasis 3 dpt following orthotopic transplant of ZMEL1-PRO or a 1:1 mixture of ZMEL1 PRO:INV (OR [95% CI]: 3.31 [1.10, 9.96]; p=0.033 by logistic regression). **e**. Proportion of fish with ZMEL1-INV caudal metastasis 3 dpt following orthotopic transplant of ZMEL1-INV or a 1:1 mixture of ZMEL1 PRO:INV (OR [95% CI]: 1.32 [0.61, 2.88]; p=0.49 by logistic regression). For **c-e**: N=5 independent experiments with PRO/MIX/INV 18/17/18, 13/15/14, 15/16/15, 12/15/15, and 15/17/16 fish per group, respectively; 231 fish total; plots show mean ± SD. **f**. Number of ZMEL1-PRO cells in Boyden Chamber assay migrating per 20X field when alone or mixed with ZMEL1-INV (p=0.042 by linear regression). **g**. Number of ZMEL1-INV cells in Boyden Chamber assay migrating per 20X field when alone or mixed with ZMEL1-PRO (p=0.91 by linear regression). For **f-g**: N=3 independent experiments for each EGFP and tdTomato labeling; plots show mean ± SD.

To test the consequences of PRO-INV interaction in a more physiological setting, we next assessed their interaction after orthotopic transplantation in adult zebrafish. We transplanted primary tumors of each population alone and as a 1:1 mixture and followed their growth and metastasis by whole-fish fluorescence microscopy. In the group with mixed primary tumors, we observed a significantly higher number of fish with polyclonal metastasis than would be expected based on the metastatic rate of each subpopulation alone if they did not interact (Figure 3b-c, Figure S4b-c). Strikingly, we also observed that the less metastatic ZMEL1-PRO population had an increased rate of caudal metastases in mixed tumors compared to when it was transplanted alone (Figure 3d, Figure S4d-h), showing that this population benefited from cell-cell interaction with the INV cells. Moreover, the more metastatic ZMEL1-INV population did not become less metastatic (Figure 3e), meaning that they did not pay a significant cost for giving this benefit to ZMEL1-PRO. This type of interaction, where one individual benefits another without paying a cost, is formally defined as cooperation (Foster, 2011). To further characterize the benefit to the ZMEL1-PRO population, we performed transplants at various mixing ratios (1:4, 4:1, and 9:1) consisting of tdTomato-expressing ZMEL1-PRO cells mixed with EGFP-expressing ZMEL1 cells (either PRO or INV) and then quantified the metastases (Figure S4i-j). This confirmed that when ZMEL1-INV cells comprise at least half of the primary tumor, the PRO subpopulation has an increased rate of metastasis, providing context to which patients may exhibit such metastatic interaction. We observed a similar cooperative interaction *in vitro* in dual-color Boyden Chamber migration assays (Figure 3f-g), confirming that ZMEL1-PRO invades better when mixed with -INV cells independently of the microenvironment. Experiments with conditioned media further suggested this interaction is mediated by direct cell-cell contact (Figure S4k-l) and not via soluble factors. The *in vivo* cooperative benefit was only evident early in metastatic dissemination (3 dpt vs 7 dpt, Figure S4g-h), indicating that this cooperation is particularly beneficial when both primary tumors and the number of disseminating tumor cells are small. Taken together, these data show that the formation of heterotypic clusters enables the collective extravasation of PRO and INV, facilitating cooperation that preserves cell state diversity in early metastatic lesions (Figure S4m) (Foster, 2011; Hauser et al., 2009).

### TFAP2 mediates the PRO/INV state and clustering

Although several molecular mechanisms have been shown to regulate the PRO and INV state in melanoma (including MITF, AXL, WNT5A and BRN2 and their up- and downstream regulatory networks (Cheng et al., 2015; Falletta et al., 2017; Fane et al., 2019; Hoek et al., 2008; Hoek et al., 2006; Pinner et al., 2009; Rambow et al., 2015; Rambow et al., 2019; Rambow et al., 2018; Shakhova et al., 2015; Tirosh et al., 2016; Verfaillie et al., 2015; Weeraratna et al., 2002; Widmer et al., 2012)), there is no known connection between these programs and the formation of tumor cell clusters. To identify the mechanism regulating clustering in the INV population, we performed motif analysis on 1 kilobase regions associated with genes differentially expressed between ZMEL1-PRO and -INV cells (Figure 4a, Supplementary Table 3). The top motif whose target genes were enriched in the PRO cells putatively binds the NFIC and TFAP2A transcription factors. One of the TFAP2 family members itself, *tfap2e*, was also one of the most differentially expressed genes between the PRO and INV cells, with its expression being over 100-fold higher in the PRO versus INV cells (Figure S5a, Supplementary Table 1). The TFAP2 family of transcription factors plays essential roles in neural crest and melanocyte cell fate during development (de Croze et al., 2011; Hoffman et al., 2007; Kaufman et al., 2016; Li and Cornell, 2007; Luo et al., 2002; Seberg et al., 2017a; Seberg et al., 2017b; Van Otterloo et al., 2010) and has been implicated as part of a regulatory network promoting the PRO state (Hoek et al., 2006; Rambow et al., 2015; Tirosh et al., 2016; Verfaillie et al., 2015). This raised the hypothesis that TFAP2 was acting as a master regulator of the clustering phenotype observed in the INV population. To test this, we performed RNA-seq of ZMEL1 cells in 3D (clustered) culture, and asked which genes were differentially expressed in 3D compared to 2D (non-clustered) conditions (Figure 4b, Figure S5b, Supplementary Tables 2,4). In both PRO and INV, we again found enrichment of a motif that binds TFAP2—specifically, TFAP2E—when looking at up- and downregulated genes together. This is consistent with the known redundancy of *tfap2a* and *tfap2e* in zebrafish (Van Otterloo et al., 2010), and highly suggestive of a role for TFAP2 in mediating clustering.

**Figure 4.**
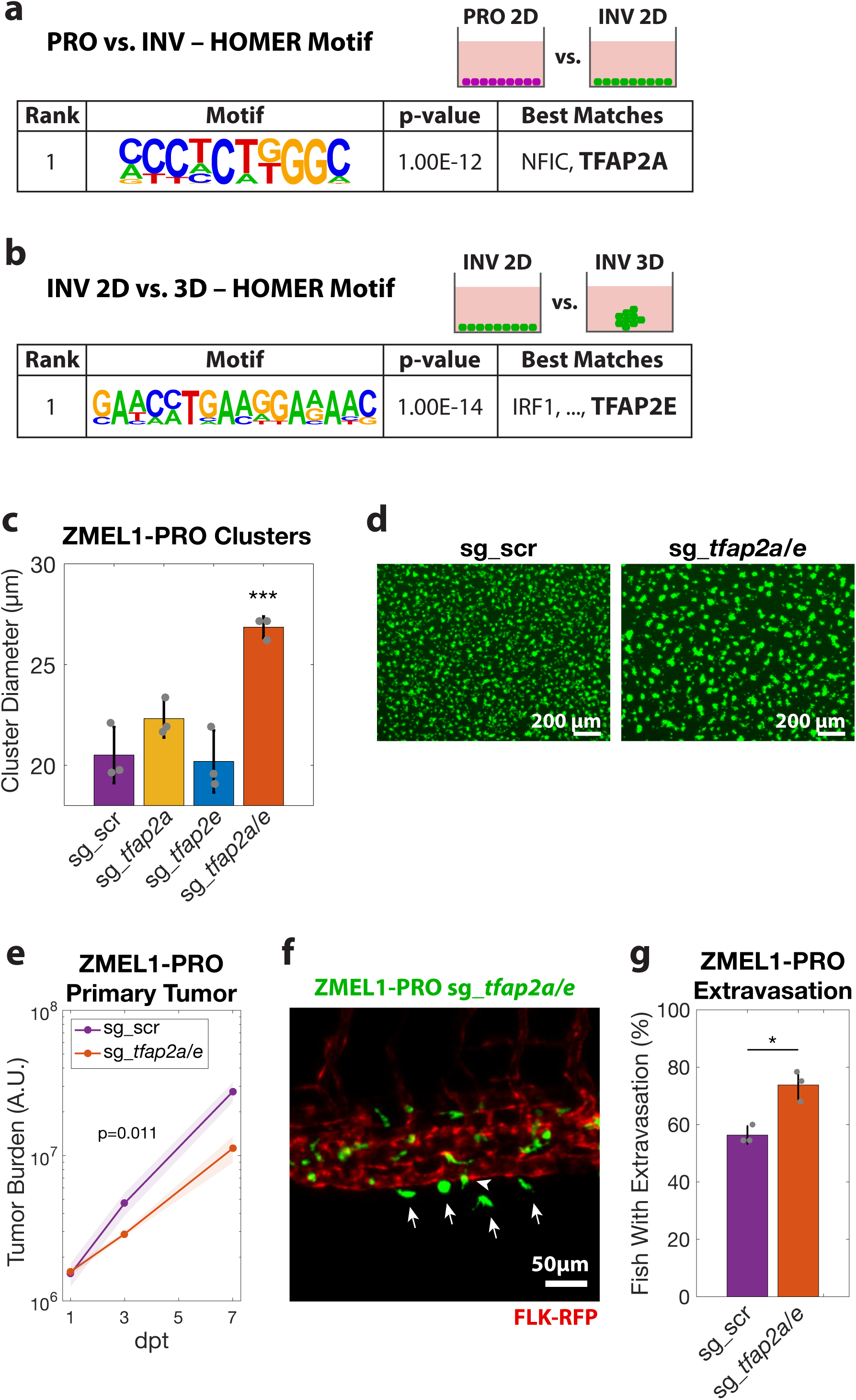
TFAP2 distinguishes PRO vs. INV state and modulates clustering and metastasis. **a**. HOMER de-novo motif analysis on genes upregulated in ZMEL1-PRO vs. -INV (log_2_ fold change cutoff ± 1.5, p_adj_ < 0.05, ±500bp of transcription start site [TSS]). **b**. HOMER de-novo motif analysis of genes differentially expressed between ZMEL1-INV in 3D (clusters) vs. 2D (no clusters) (log_2_ fold change cutoff ± 1.5, p_adj_ < 0.05, ±500bp of TSS). **c**. Cluster size after 2 days in ZMEL1-PRO with CRISPR-Cas9 inactivation of *tfap2a* and *tfap2e* alone or in combination (sg_*tfap2a/e*) versus control (sg_scr) (p-values by linear regression; N=3 independent experiments). **d**. Representative images of clusters formed after 2 days from ZMEL1-PRO with sg_scr vs. sg_*tfap2a/e*. **e**. Growth of ZMEL1-PRO orthotopic primary tumors with sg_scr vs. sg_*tfap2a/e* (p=0.011 by linear regression; N=3 independent experiments with sg_scr/sg_tfap2a/e 24/22, 22/22, 24/24 fish per group, respectively; n=138 fish total). **f**. Representative image of extravasated (arrows) and partially extravasated (arrow-head) ZMEL1-PRO cells with sg_*tfap2a*/*e* following intravenous transplant in *casper* fish with FLK-RFP transgene labeling the vasculature. **g**. Proportion of larval fish intravenously transplanted with ZMEL1-PRO with sg_scr or sg_*tfap2a/e* with extravasated cells at 1 dpt, as quantified from confocal time lapse microscopy (OR [95% CI]: 2.20 [1.05, 4.61]; p=0.038 by logistic regression; N=3 independent experiments with sg_scr/sg_tfap2a/e 20/20, 22/23, and 22/22 fish per group, respectively; n=129 fish total).

We next sought to test whether TFAP2 plays a functional role in melanoma cluster formation and metastasis. We performed CRISPR/Cas9 deletion of *tfap2a* and *tfap2e* in ZMEL1-PRO, which typically forms poor clusters, and found a significant increase in clustering only in the context of *tfap2a/e* double knockout (Figure 4c-d, Figure S5c-e). We also found that the *tfap2a/e* knockout compared with a non-targeting control had a small but reproducible decrease in cell proliferation, along with an increase in the persistence of migration (Figure S5f-i), consistent with the phenotype differences between the INV and PRO populations. We next wanted to determine whether this phenotypic switch mediated by TFAP2 translated to an *in vivo* effect on metastasis. We orthotopically transplanted control or *tfap2a/e* knockout cells into adult *casper* fish and measured both primary tumor growth and metastatic dissemination. The *tfap2a/e* knockout cells formed primary tumors that grew significantly slower than controls (Figure 4e), which was expected from their slower *in vitro* proliferation. Despite this decrease in primary tumor growth, we found similar rates of overall and caudal metastasis, suggesting that loss of *tfap2a/e* induces a higher proportion of cells to metastasize (Figure S5j-k). To test this idea more directly, we assessed the effect of *tfap2a/e* on metastasis in a proliferation-independent assay by intravenous transplant. Time lapse confocal microscopy revealed that loss of *tfap2a/e* led to metastatic extravasation in a significantly higher proportion of fish (Figure 4f-g), consistent with a report that TFAP2A overexpression in human cells slows metastatic spread (Huang et al., 1998). Taken together, these data suggest that TFAP2 is not only a major regulator of the PRO vs. INV cell state, but that it also controls tumor cell clustering and regulates metastasis via an effect on extravasation.

### TFAP2 correlates with clustering in human melanoma

We next wanted to determine whether the effects of TFAP2 we observed in the zebrafish were conserved in human melanoma. TFAP2A is a member of several gene expression profiles describing the proliferative state (Rambow et al., 2015; Tirosh et al., 2016; Verfaillie et al., 2015), including the Hoek et al. set (Hoek et al., 2006), and is critical for melanoma cell proliferation (Figure S6a). Consistent with this, increased expression in primary tumors of either TFAP2A or genes associated with the PRO state is associated with worse clinical outcomes in two large independent clinical cohorts (Figure S6b-g), likely reflecting the known prognostic effects of mitotic rate and primary tumor size in melanoma TNM staging (Gershenwald et al., 2017; Thompson et al., 2011). We next examined TFAP2A expression in a panel of 56 melanoma cell lines (CCLE) (Ghandi et al., 2019) and 472 clinical melanoma samples (TCGA) (Cancer Genome Atlas Network, 2015), and asked how this correlated with their PRO/INV signatures defined by Hoek et al. (Hoek et al., 2006). In both cohorts, we confirmed that the PRO and INV states were strongly anti-correlated. Samples with higher TFAP2A expression exhibited a more PRO gene signature, and conversely, samples with lower TFAP2A expression exhibited a more INV signature (Figure 5a, Figure S6h). Further, we asked whether the association between TFAP2A expression and the PRO/INV signatures was maintained at the level of single cells. We analyzed available single cell RNA-seq data across a panel of 23 human melanoma patients (Jerby-Arnon et al., 2018), and found a similar relationship: individual cells with high TFAP2A tend to have a higher PRO score, whereas cells with low TFAP2A tend to have a higher INV score (Figure S6i).

**Figure 5.**
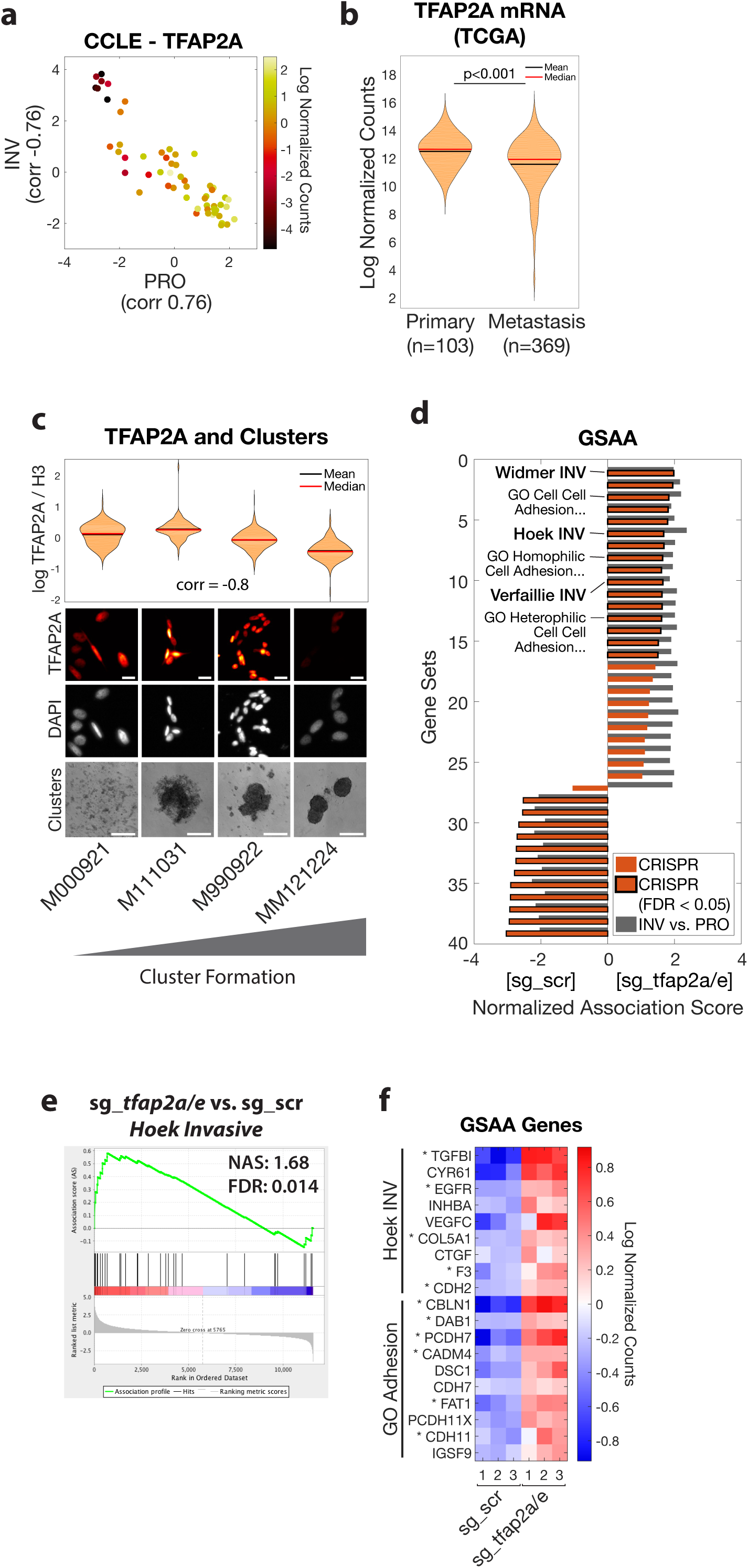
TFAP2 correlates with clustering in human melanoma and regulates genes associated with metastasis and cell-cell adhesion. **a**. Human melanoma cell lines in the Cancer Cell Line Encyclopedia (CCLE, n=56) plotted as PRO and INV scores (Hoek et al., 2006) calculated from RNA-seq and colored according to TFAP2A mRNA expression. Pearson correlation coefficients between TFAP2A and PRO/INV scores are shown on axes. **b**. TFAP2A mRNA expression in The Cancer Genome Atlas (TCGA) melanoma (SKCM) cohort comparing primary tumors and metastases (p<0.001 by Wilcoxon rank sum test with Bonferroni correction). **c**. Low-passage human melanoma cell lines ranked by increased cluster forming ability (left to right) with TFAP2A expression quantified by immunofluorescence (plot and top; Spearman correlation shown; scale bar 20 μm) and clustering (bottom, scale bar 500 μm). **d**. GSAA was run using gene sets and GO gene sets with FDR < 0.05 from INV vs. PRO RNA-seq (n=39 gene sets; cyan points in Figures 1d and 2a). Bars show Normalized Association Score (NAS) for CRISPR (ZMEL1-PRO sg_*tfap2a/e* vs. sg_scr) and INV vs. PRO for each gene set, with black outline representing FDR<0.05 for CRISPR experiment. **e**. Plot of Hoek et al. (Hoek et al., 2006) INV signature by GSAA for ZMEL1-PRO sg_*tfap2a/e* vs. sg_scr RNA-seq. **f**. Heatmap of top genes in Hoek INV and GO Adhesion gene sets that are differentially expressed between ZMEL1-PRO sg_*tfap2a/e* and sg_scr (log_2_ fold change cutoff ± 0.5, p_adj_ < 0.05). Asterisks (*) indicate genes with associated TFAP2A CUT&RUN peaks. Human ortholog gene names are used for clarity (see Figure S7a for zebrafish gene names).

Within the TCGA dataset, tumor samples collected from primary sites had higher levels of TFAP2A compared to metastatic lesions despite similar expression of pan-melanoma markers (Figure 5b, Figure S6j-k), in agreement with a prior report (Tellez et al., 2007). Further, in the two patients for which paired primary and metastatic samples were available, TFAP2A expression was lower in the metastatic lesion. A direct measurement of the relative ratio of TFAP2^HI^ to TFAP2^LO^ cells in the tumors, and its correlation with patient prognosis will await future longitudinal prospective single cell analysis. Next, we examined TFAP2A expression in the panel of human melanoma cell lines used in Figure 2e and found that cluster-forming lines had lower TFAP2A expression than non-clustering lines (Figure S6l-m). To test this association across cells that better preserve the heterogeneity observed clinically in melanoma, we examined a panel of four short-term human melanoma cultures (Raaijmakers et al., 2015). Cluster formation correlated strongly with lower expression of TFAP2A (Figure 5c), consistent with our observation that TFAP2 loss drives melanoma clustering. Collectively, our data confirm that the association we had discovered in zebrafish—between TFAP2, the PRO/INV state and tumor cell clustering—also occurs in human melanoma.

### TFAP2 regulates genes associated with metastasis and cell-cell adhesion

To gain further insight into the mechanism by which TFAP2 regulates melanoma phenotypes, we performed RNA-seq of the *tfap2a/e* knockout cells versus controls. We first validated that the *tfap2a/e* knockout recapitulated the observed differences between ZMEL1-PRO and -INV by performing gene set association analysis (GSAA) using the gene sets that had passed false discovery cutoff (FDR < 0.05) in our ZMEL1-INV vs. -PRO RNA-seq analysis. We observed a high concordance in the top dysregulated pathways—including multiple INV and GO adhesion gene sets associated with TFAP2 loss—confirming that TFAP2 regulates pathways distinguishing ZMEL1-PRO and -INV (Figure 5d-e, Supplementary Tables 5,6,7). Specific genes upregulated upon TFAP2 loss and associated with either the INV state or adhesion include several with known functions in melanoma metastasis (Figure 5f, Figure S7a; e.g. TGFBI (Lauden et al., 2014), VEGFC (Streit and Detmar, 2003), CTGF (Finger et al., 2014), and CDH2 (Mrozik et al., 2018)). In order to understand the mechanism by which TFAP2 regulates PRO/INV state and cell-cell adhesion, we performed TFAP2A CUT&RUN (Cleavage Under Targets and Release Using Nuclease) in SKMEL28 cells, the human cell line with the highest expression of PRO-state genes out of those we characterized. This allowed us to understand the genes bound by TFAP2A in melanoma (Rambow et al., 2015; Seberg et al., 2017b). Consistent with the known roles of TFAP2 as both a transcriptional activator and repressor (Ren and Liao, 2001; Seberg et al., 2017b), we observed significant enrichment for TFAP2A peaks in genes that are upregulated upon *tfap2a/e* knockout in ZMEL1-PRO cells (Figure 5f asterisks, Figure S7b-d; e.g. TGFBI, CDH2), suggesting it acts as a repressor of those loci. We did not observe evidence of a stress response resulting from *tfap2a/e* knockout (Supplementary Table 7), lending further support to a model of direct regulation by TFAP2. Taken together, these data highlight the direct and pleiotropic effects of TFAP2 loss on metastatic spread, further confirming a role for TFAP2 in cell state and suggesting downstream mediators.

### Longitudinal single-cell RNA-seq reveals stability of PRO but not INV state

The above data suggest a model in which PRO and INV cell clusters, regulated by TFAP2, form the unit of initial metastatic seeding. However, once seeding has occurred, it is still possible that either of these cell states can undergo phenotype switching and contribute to metastatic outgrowth. This possibility was suggested by our finding that metastases tend to become dominated by PRO cells over time (Figure 6a-b). To test this more formally, we conducted a large-scale longitudinal analysis of cell state at the single cell level, interrogating the effects of cell-cell interaction, tumor formation, and metastasis. We performed single-cell RNA-seq on over 40,000 ZMEL1 cells from both the PRO and INV cell states across four different conditions: (1) *in vitro* individual culture; (2) *in vitro* co-culture; (3) *in vivo* primary tumors; and (4) *in vivo* metastatic lesions (Figure 6a). Strikingly, ZMEL1-PRO and -INV subpopulations were highly pure at baseline and remained largely discrete throughout all conditions (Figure 6c). In order to quantify the stability of the two populations, we calculated PRO and INV scores for each cell based on gene sets derived from ZMEL1 bulk RNA-seq, and trained a classifier based on *in vitro* individual culture samples (Figure 6d-e). Consistent with our results from conditioned media experiments (Figure S4k-l), we observed very little effect of co-culture upon the transcriptomes of ZMEL1-PRO and -INV cells, with both populations remaining more than 99% pure. Strikingly, a fraction of INV cells in tumors, especially from the metastases, upregulated PRO-state genes, increasingly occupying a PRO/INV double-positive state. This is in contrast to PRO cells, which remained stable in the PRO state. Further validating a role for TFAP2 as a master regulator of melanoma cell state, we found that ZMEL1-INV cells that gained a PRO-like gene expression program also reactivated *tfap2e* (Figure 6f, Figure S8a). Overall, these data support a model of cooperation whereby clusters comprised of distinct PRO and INV subpopulations promote co-metastatic seeding, and metastatic outgrowth is increasingly dominated by PRO-like cells.

**Figure 6.**
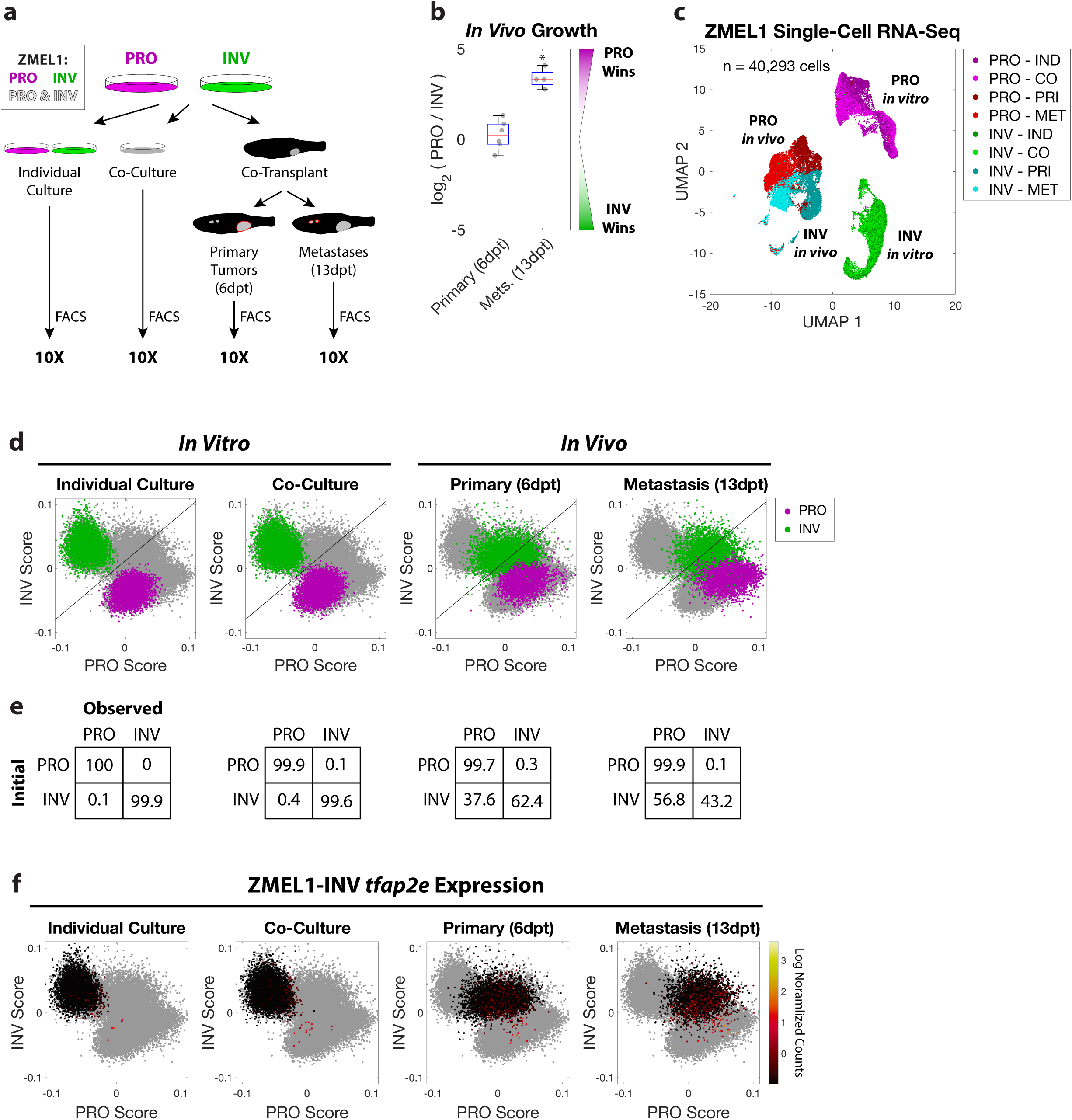
Longitudinal single-cell RNA-seq reveals stability of PRO but not INV state. **a**. Schematic of experiment. Prior to flow cytometry and 10X single-cell RNA-seq, ZMEL1-PRO and -INV cells were either grown *in vitro* (individual or co-culture) or isolated from zebrafish orthotopically transplanted with a 1:1 mixture of the two subpopulations (primary tumors or metastases). **b**. Relative number of ZMEL1-PRO and -INV cells isolated and quantified by flow cytometry from primary tumors and metastases of fish transplanted with a 1:1 mixture of ZMEL1-PRO and -INV (primary tumors from n=6 fish; metastases from n=4 fish; p= 0.51 and p=0.031, respectively, by one-sample two-sided t-test with Bonferroni correction). **c**. Uniform Manifold Approximation and Projection (UMAP) (McInnes et al., 2018) dimensionality reduction of 40,293 ZMEL1 cells sequenced as in (a). Individual culture (IND); co-culture (CO); primary tumors (PRI); metastases (MET). **d**. PRO and INV scores based on ZMEL1 bulk RNA-seq are plotted for all cells in gray. ZMEL1-PRO (purple) and ZMEL1-INV (green) for the indicated condition are colored. Diagonal line represents the classifier used in (e). **e**. Confusion matrices comparing initial cell identity with observed cell classification based on a linear classifier trained on *in vitro* individual culture samples. **f**. ZMEL1-INV cells plotted as in (d) colored according to *tfap2e* mRNA expression reveal re-activation of *tfap2e* upon metastatic dissemination.

## DISCUSSION

Both individual and collective mechanisms of metastasis can occur in melanoma (Long et al., 2016) and other cancers (Pearson, 2019; Reichert et al., 2018). Phenotype switching between PRO and INV states has long been postulated to be a mechanism for individual seeding of metastasis in melanoma (Hoek et al., 2008; Kim et al., 2017; Pinner et al., 2009; Vandamme and Berx, 2014). Separately, circulating tumor cell (CTC) clusters, a mode of collective metastasis, have been shown to have increased metastatic potential (Aceto et al., 2014; Cheung et al., 2016), and patients with detected CTC clusters have worse clinical outcomes (Giuliano et al., 2018; Long et al., 2016). Cooperation has previously been reported both in epithelial cancers (Celià-Terrassa et al., 2012; Neelakantan et al., 2017; Tsuji et al., 2009) and between melanoma PRO and INV states in the context of primary tumor collective cell invasion (Chapman et al., 2014) and metastatic tropism (Rowling et al., 2020), but the mechanisms that explain the relationship between the PRO/INV states and cooperative metastasis have remained unknown. We provide for the first time a clear mechanism that explains how these two subpopulations, which coexist in the primary tumor, cooperate in metastasis formation. We find that PRO and INV cells form heterotypic clusters which are controlled by the neural crest transcription factor TFAP2. It is likely that in individual patients, either individual or collective migration may predominate, consistent with the observation that a subset of patients have clusters comprised of cells with variable MITF levels (Khoja et al., 2014) and exhibit polyclonal metastatic seeding (Rabbie et al., 2019; Sanborn et al., 2015). While the phenotype switching model predicts dynamic switching of individual cells between PRO and INV states (analogous to an epithelial-to-mesenchymal transition) as a necessary feature of individual metastasis (Hoek et al., 2008; Kim et al., 2017; Pinner et al., 2009; Vandamme and Berx, 2014), our finding that PRO and INV can cooperate while remaining as distinct phenotypic populations suggests that tumors can preserve diversity during initial metastatic seeding without the need for cell state switching on a rapid time scale. However, our single cell analysis of metastatic outgrowth demonstrates that INV cells, once they arrive, can still switch to a double-positive PRO/INV state, indicating that phenotype switching in the INV to PRO direction may be operative after initial seeding.

Our data demonstrate that cell cluster formation driven by TFAP2 loss is a pro-metastatic feature of INV cells, with pleiotropic increases in cell-cell adhesion and cell clustering enabling cooperation with PRO cells. Further, we demonstrate the functional role of TFAP2 in regulating cell state and clustering. How TFAP2 itself is regulated in this context, however, remains an open question. DNA methylation has been linked to expression of PRO/INV genes (Verfaillie et al., 2015) and to TFAP2A expression (Hallberg et al., 2014; Zeng et al., 2013); however, further work is required to fully elucidate these relationships. This mechanism is consistent with the recent report that breast cancer epigenetic state and CTC cluster formation are tightly linked (Gkountela et al., 2019), and suggests that clusters may act to potentiate an already more metastatic cell population. Given that the INV state in melanoma is also associated with increased resistance to targeted therapy (Konieczkowski et al., 2014; Muller et al., 2014; Verfaillie et al., 2015), pharmacologic disruption of CTC clusters could be an attractive target to slow metastasis and decrease the distant spread of drug-resistant cells.

## ACKNOWLEDGEMENTS

This work was supported by an NIH Research Program Grant under award number R01CA229215 to J.B.X. and R.M.W. N.R.C. was supported by the Kirschstein-NRSA predoctoral fellowship (F30) from the NIH under award number F30CA220954, by a research grant from the Melanoma Research Foundation, and by a Medical Scientist Training Program grant from the NIH under award number T32GM007739. This work was also supported by an NIH Research Program Grant under award number R01AR062547 to R.A.C. The Leeds Melanoma cohort was funded by Cancer Research UK grants C588/A19167, C8216/A6129, and C588/A10721 and NIH grant R01CA83115. We thank the University Research Priority Program (URPP) of the University of Zurich for access to the melanoma biobank and early passage cultures. The results published here are in part based upon data generated by the TCGA Research Network: https://www.cancer.gov/tcga.

## AUTHOR CONTRIBUTIONS

N.R.C., J.B.X. and R.M.W. designed the study and wrote the manuscript, on which all authors commented. N.R.C., A.R., M.Z., T.-H.H., E.M., M.T., and M.H. performed experiments. C.K. and R.A.C. performed and analyzed CUT&RUN experiments. N.R.C. analyzed *in vitro, in vivo*, and RNA-seq data. N.R.C., A.R., M.B., and I.Y. analyzed single-cell RNA-seq data. S.H. developed methods for *in vivo* imaging and analysis. M.D. developed methods for *in vitro* cell tracking and image analysis. L.F. and M.P.L. generated low-passage patient-derived cell lines. M.G., J.N., J.N.-B., M.R.M., P.C., D.J.A. and R.R. analyzed clinical data.

---

**Figure S1.**
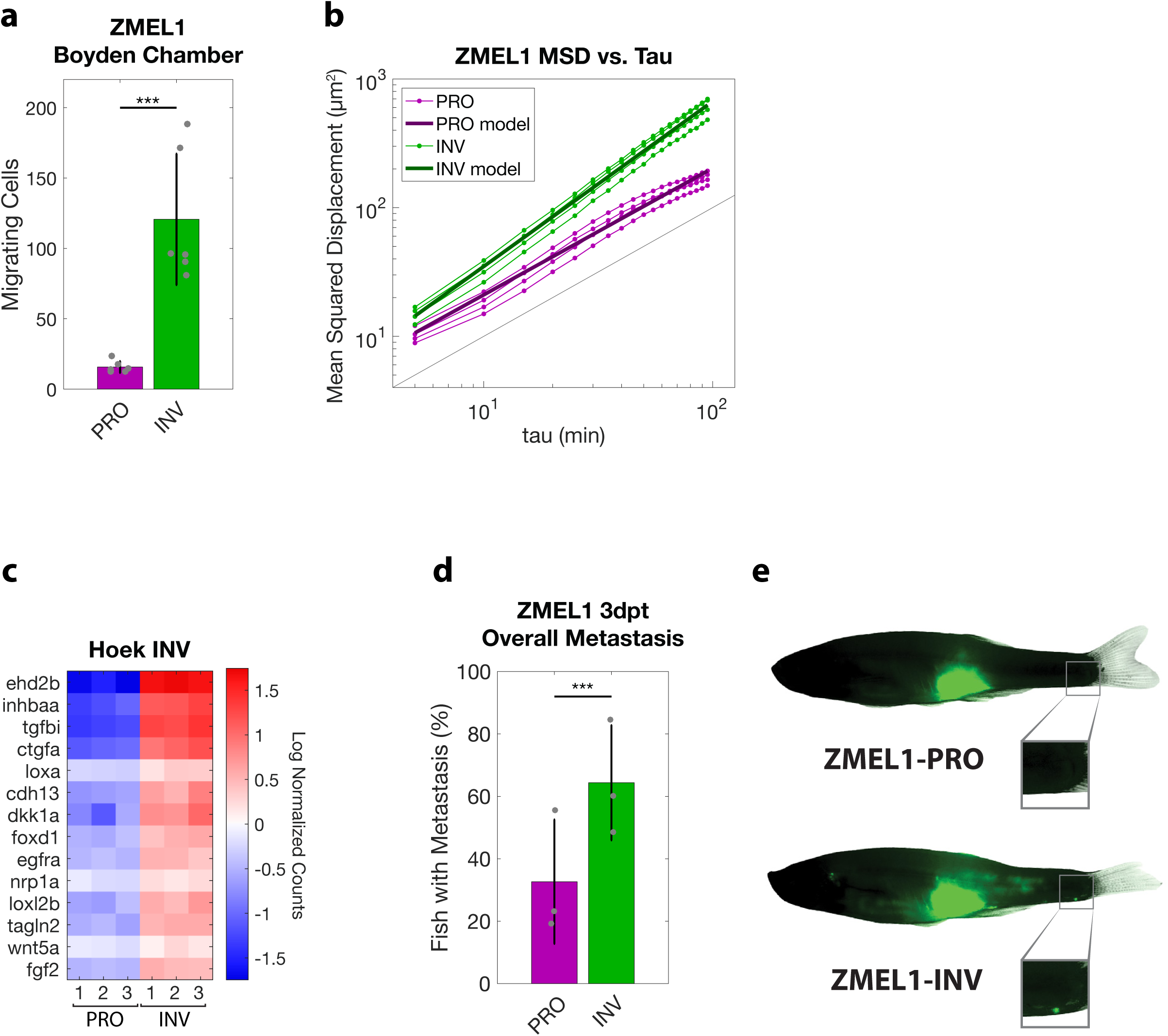
(Related to Figure 1) **a**. Number of ZMEL1 cells migrating in Boyden Chamber assay (p<0.001 by linear regression, N=3 independent experiments for each of 2 fluorophores). **b**. Log-log plot of mean squared displacement (MSD) vs. lag time (tau) over the range of 5≤tau<100 minutes with model fits overlaid (N=4 independent experiments, see Methods for details). The slope (α) provides quantification of the persistence of motility, where a cell moving randomly will have α=1, and a cell moving along a straight line will have α=2 (Gorelik and Gautreau, 2014). Black line with α=1 is shown for comparison. **c**. Heatmap of genes in the Hoek INV signature that are differentially expressed in ZMEL1-INV vs. -PRO (log_2_ fold change cutoff ± 1.5, p_adj_<0.05). As in Figure 1e, but with zebrafish gene names. **d**. Quantification of overall distant metastases seeded by ZMEL1 populations at 3 dpt (OR [95% CI]: 4.35 [1.87, 10.11]; p<0.001 by logistic regression; N=3 independent experiments with PRO/INV 9/10, 31/33, and 13/13 fish per group, respectively; n=109 fish total; plot shows mean ± SD). **e**. Representative images of ZMEL1-PRO and -INV tumors and distant metastases (e.g. to caudal region [box]) at 3 days post-transplant (3dpt).

**Figure S2.**
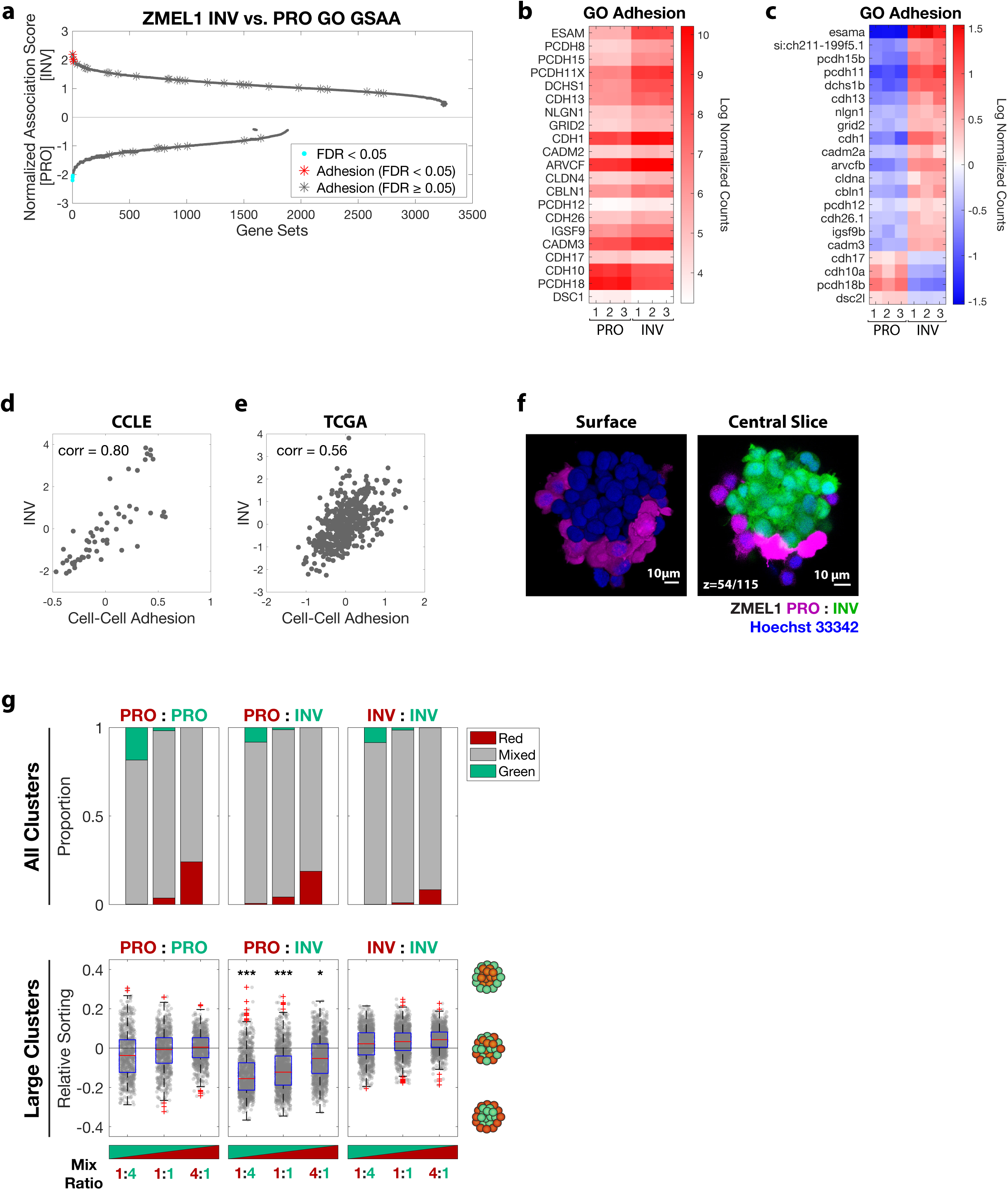
(Related to Figure 2) **a**. Dual waterfall plot of all gene sets from GO analysis. Adhesion GO gene sets are indicated with an asterisk and colored according to false discovery rate (FDR). **b-c**. Heatmap of genes in adhesion GO gene sets (FDR < 0.05, n=3) that are differentially expressed between ZMEL1-PRO and -INV (log_2_ fold change cutoff ± 1.5, p_adj_ < 0.05). As in Figure 2b, but with (b) absolute expression data, and (c) zebrafish gene names. **d-e**. Human melanoma samples from (d) The Cancer Cell Line Encyclopedia (CCLE, n=56) and (e) The Cancer Genome Atlas (TCGA) melanoma (SKCM, n=472) cohort are plotted as INV (Hoek et al., 2006) versus Cell-Cell Adhesion scores calculated from RNA-seq. Pearson correlation coefficient between scores is shown. **f**. (left) 3D opacity rendering and (right) central slice of confocal imaging through co-cluster of ZMEL1-PRO and ZMEL1-INV. **g**. Quantification of (top) mixing and (bottom) sorting of heterotypic ZMEL1 clusters as in Figure 2f. Red (tdTomato) and green (EGFP) labeled ZMEL1 cells were mixed PRO:PRO, PRO:INV, and INV:INV at indicated ratios. (top) Quantification of all clusters revealed that nearly all clusters mix, regardless of cell type. (bottom) Spatial sorting was significantly enriched in PRO:INV clusters compared to PRO:PRO and INV:INV controls (p<0.001, p<0.001, and p=0.038 for 1:4, 1:1, and 4:1, respectively, by one-way ANOVA on mean of each replicate versus respective PRO:PRO and INV:INV controls [greater of two p-values reported]).

**Figure S3.**
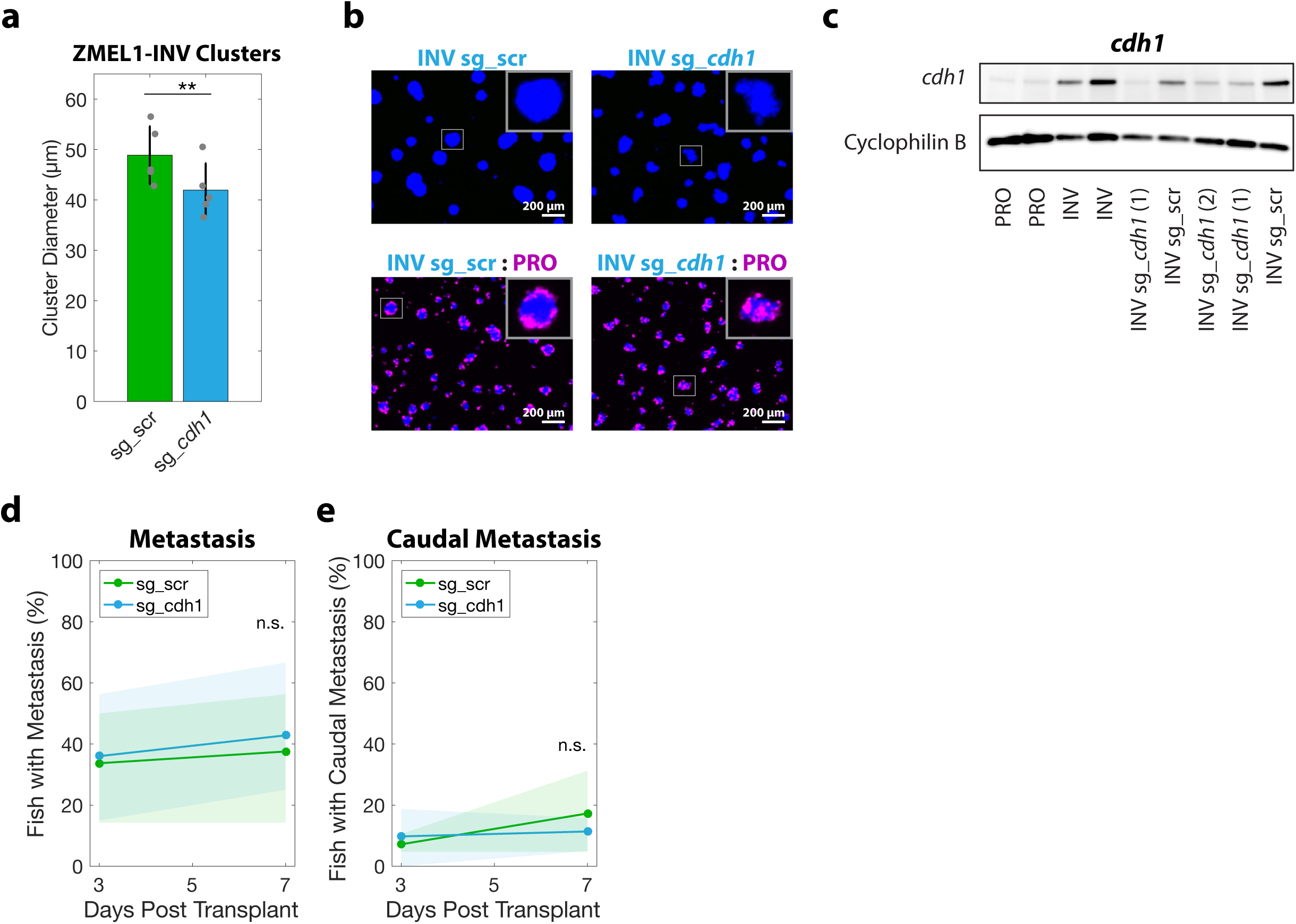
(Related to Figure 2) **a**. Cluster size after 2 days in ZMEL1-INV with CRISPR deletion of *cdh1* (sg_*cdh1*) versus control (sg_scr) (p=0.0016, two-tailed paired t-test, N=5 independent experiments). **b**. (top) Representative images of effect of sg_*cdh1* versus sg_scr on cluster size at 3 days. (bottom) Representative images of mixing effects of sg_*cdh1* versus sg_scr. Control (sg_scr) ZMEL1-INV cells mixed with ZMEL1-PRO show clear spatial sorting, whereas sg_*cdh1* ZMEL1-INV cells mixed with ZMEL1-PRO demonstrate decreased sorting. **c**. Western blot confirmed two different sg_*cdh1*’s decreased Cdh1 protein expression in ZMEL1-INV to a level comparable to ZMEL1-PRO. sg_*cdh1* (1) was utilized for all phenotypic experiments. **d-e**. ZMEL1-INV orthotopic primary tumors with sg_*cdh1* do not seed (d) distant metastases and (e) caudal metastases in a significantly different percentage of zebrafish than with sg_scr control (p=0.56 and p=0.44, respectively, at 7 dpt by logistic regression; N=3 independent experiments with sg_scr/sg_cdh1 16/15, 21/20, 19/19 fish per group, respectively; n=110 fish total).

**Figure S4.**
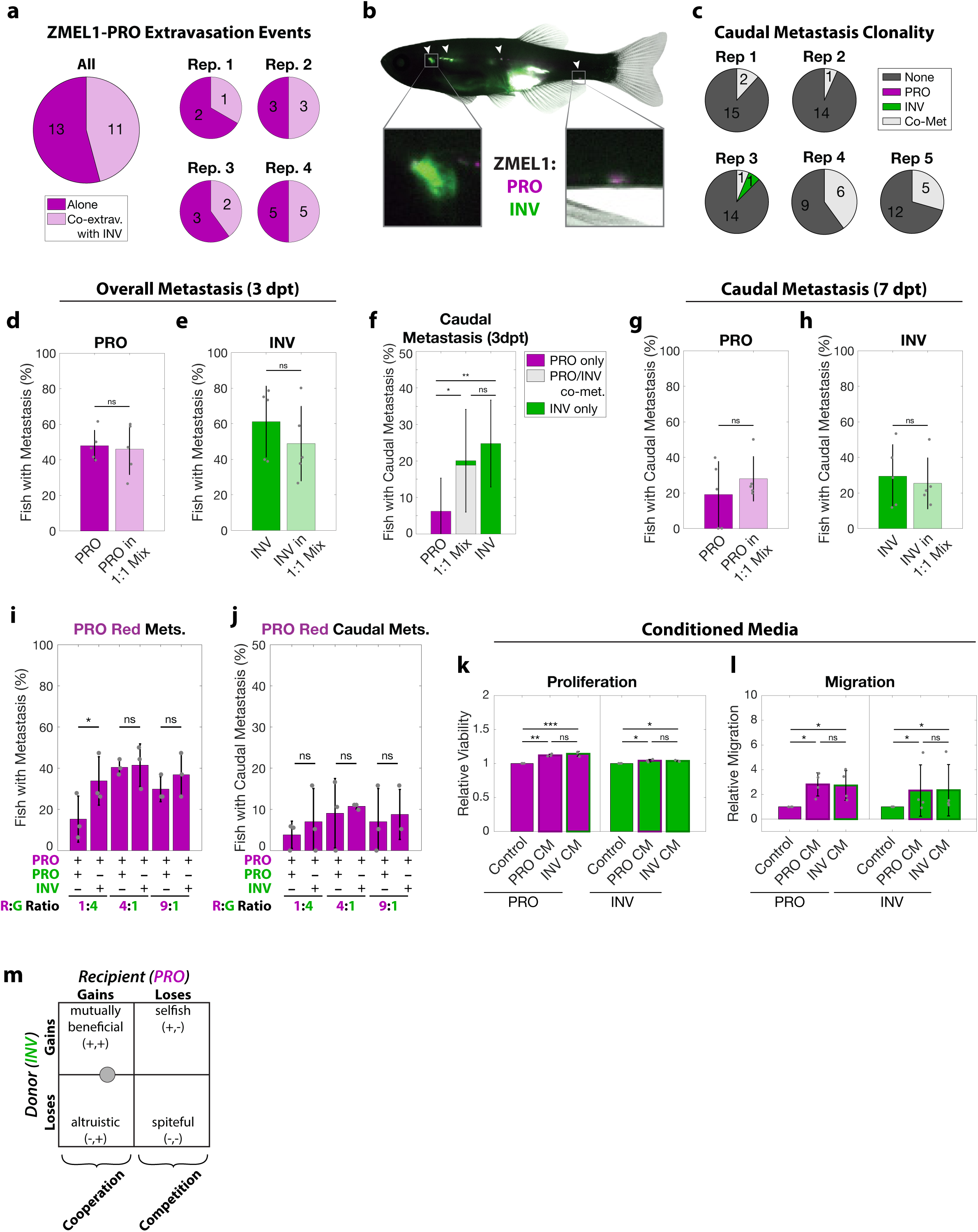
(Related to Figure 3) **a**. Number of observed extravasation events of ZMEL1-PRO either alone or in the form of co-extravasation with ZMEL1-INV following intravenous transplant (p=0.18 by two-tailed paired t-test, N=4 independent experiments with 14, 15, 22, and 22 fish each; n=73 fish total). **b**. Adult zebrafish with orthotopic transplants of 1:1 mixtures of ZMEL1-PRO and -INV seed polyclonal metastases (arrowheads), including to the kidney and caudal regions (left and right boxes, respectively). **c**. Number of fish co-transplanted with a 1:1 mixture of ZMEL1-PRO and -INV that have no caudal metastases (None), caudal metastases comprised of exclusively PRO or INV, or caudal metastases formed by co-metastasis (Co-Met) of both cell types (N=5 independent experiments with 17, 15, 16, 15, and 17 fish each; 80 fish total; p<0.001 by Mantel-Haenszel’s test for null hypothesis of no interaction; as in Figure 3c for each independent experiment). **d**. ZMEL1-PRO and **e**. ZMEL1-INV showed similar levels of overall metastasis at 3 dpt in mixed tumors compared to when each was transplanted alone (p=0.83 and p=0.13, respectively, by logistic regression). **f**. Proportion of fish with caudal ZMEL1-PRO only, ZMEL1-INV only, or PRO/INV co-metastasis (co-met) at 3dpt (PRO vs. INV: p=0.0054; “PRO only” in PRO vs. 1:1 mix: p=0.033; “INV only” in INV vs. 1:1 mix: p=0.49; all by logistic regression). Alternate presentation of data in Figure 3d-e. **g-h**. (g) ZMEL1-PRO and (h) ZMEL1-INV showed similar levels of caudal metastasis at 7 dpt in mixed tumors compared to when each was transplanted alone (p=0.54 and p=0.63, respectively, by logistic regression). For **b-h**: N=5 independent experiments with PRO/MIX/INV 18/17/18, 13/15/14, 15/16/15, 12/15/15, and 15/17/16 fish per group, respectively; 231 fish total; plots show mean ± SD). **i-j**. Adult zebrafish were orthotopically transplanted with an equivalent final concentration of tdTomato-labeled ZMEL1-PRO mixed at a 1:4, 4:1, or 9:1 ratio with EGFP-labeled ZMEL1-PRO or ZMEL1-INV. tdTomato-labeled ZMEL1-PRO (i) overall metastases and (j) caudal metastases were quantified 3dpt to measure the impact of cooperation with ZMEL1-INV. ZMEL1-PRO metastasized more when co-transplanted with ZMEL1-INV than with ZMEL1-PRO in Red:Green 1:4 transplants (OR [95% CI]: 2.78 [1.09, 7.10]; p=0.032 by logistic regression). A total of N=3 independent variable ratio mixing experiments were performed, each with at least 16 fish per group (n=108/113/113 fish per replicate; n=334 fish total). **k-l**. Proliferation (k) and Boyden chamber migration (l) were quantified in ZMEL1-PRO and -INV cells with or without conditioned media (CM) from ZMEL1-PRO and -INV cells (p-values by linear regression; N=3 independent experiments for proliferation; N=4 independent experiments for migration). **m**. The interaction between INV (donor) and PRO (recipient) cells can be schematically represented as a donor-recipient interaction (Hauser et al., 2009) falling into a regime of cooperation.

**Figure S5.**
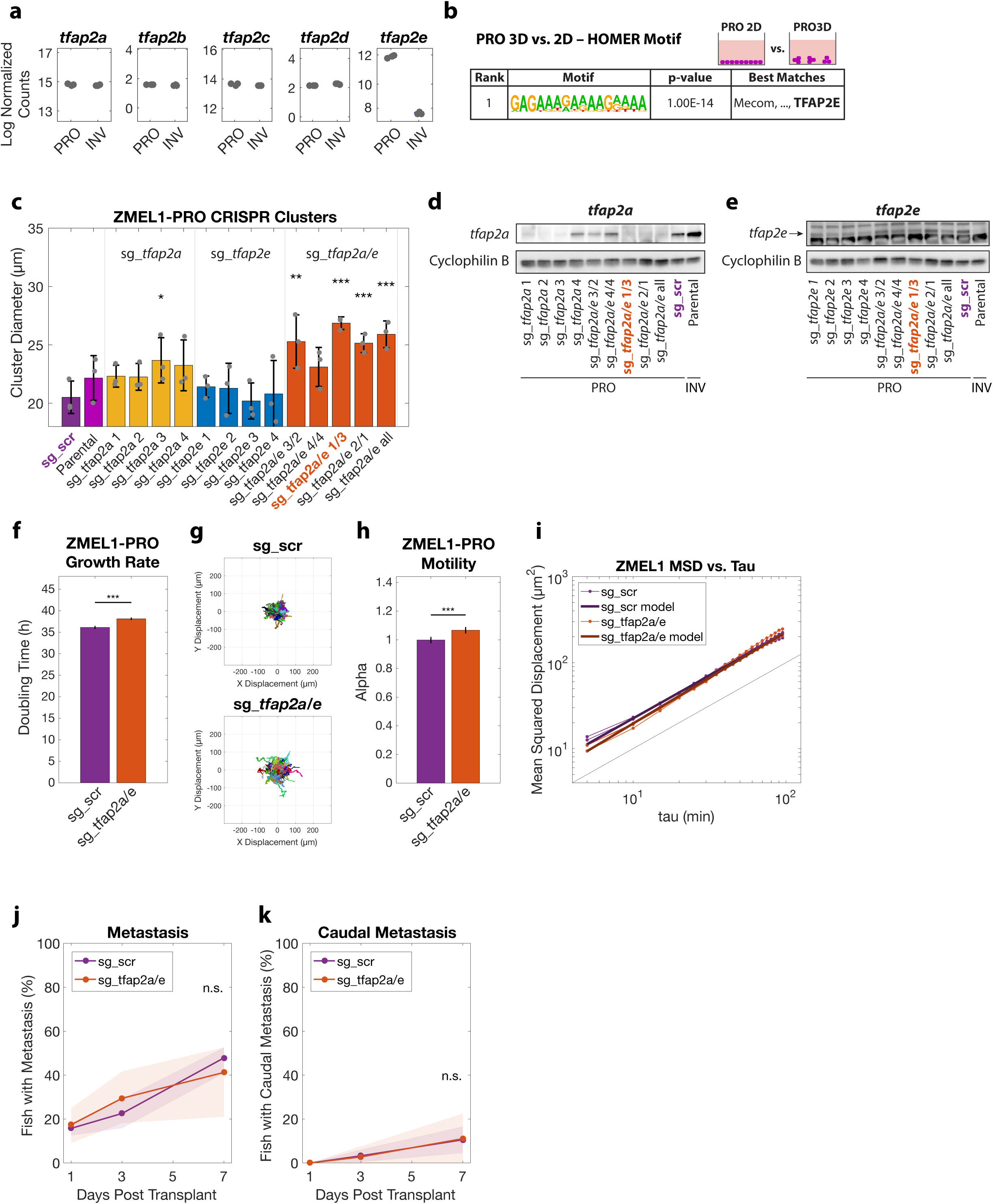
(Related to Figure 4) **a**. Boxplots of *tfap2a-e* expression from RNA-seq of ZMEL1-PRO and -INV. **b**. HOMER de-novo motif analysis of genes differentially expressed between ZMEL1-PRO in 3D (clusters) vs. 2D (no clusters) (log_2_ fold change cutoff ± 1.5, p_adj_ < 0.05, ±500bp of transcription start site [TSS]). **c**. Cluster size in ZMEL1-PRO with CRISPR-Cas9 inactivation of *tfap2a* or *tfap2e* alone and in combination (p-values by linear regression; N=3 independent experiments). sgRNAs highlighted in purple (sg_scr) and orange (sg_*tfap2a/e* 1/3) were used for further experiments (Figure 4 and Figure S5,7). **d-e**. Western blot confirmation of CRISPR inactivation of (d) *tfap2a* and (e) *tfap2e* with each sgRNA or combination of sgRNAs. **f-i**. Tracking of individual cells by time-lapse microscopy (N=3 independent experiments). **f**. ZMEL1-PRO with sg_*tfap2a/e* has slowed growth versus sg_scr (p<0.001 by linear regression, model estimates ± 95% CI shown). **g**. Representative displacements of 500 tracks per sgRNA. **h**. Model estimates ± 95% CI for alpha, the slope of the log-log plot of mean squared displacement vs. lag time (tau) for each ZMEL1-PRO sg_*tfap2a/e* and sg_scr (p<0.001 by linear regression). Larger alpha indicates more persistent motion. **i**. Log-log plot of mean squared displacement (MSD) vs. lag time (tau) over the range of 5≤tau<100 minutes with model fits overlaid (see Methods for details). The slope (α) provides quantification of the persistence of motility, where a cell moving randomly will have α=1, and a cell moving along a straight line will have α=2 (Gorelik and Gautreau, 2014). Black line with α=1 is shown for comparison. **j-k**. ZMEL1-PRO orthotopic primary tumors with sg_*tfap2a/e* do not seed (j) distant metastases and (k) caudal metastases in a significantly different proportion of zebrafish than with sg_scr control (p=0.44 and p=0.90, respectively, at 7 dpt by logistic regression; N=3 independent experiments with sg_scr/sg_tfap2a/e 24/22, 22/22, 24/24 fish per group, respectively; n=138 fish total).

**Figure S6.**
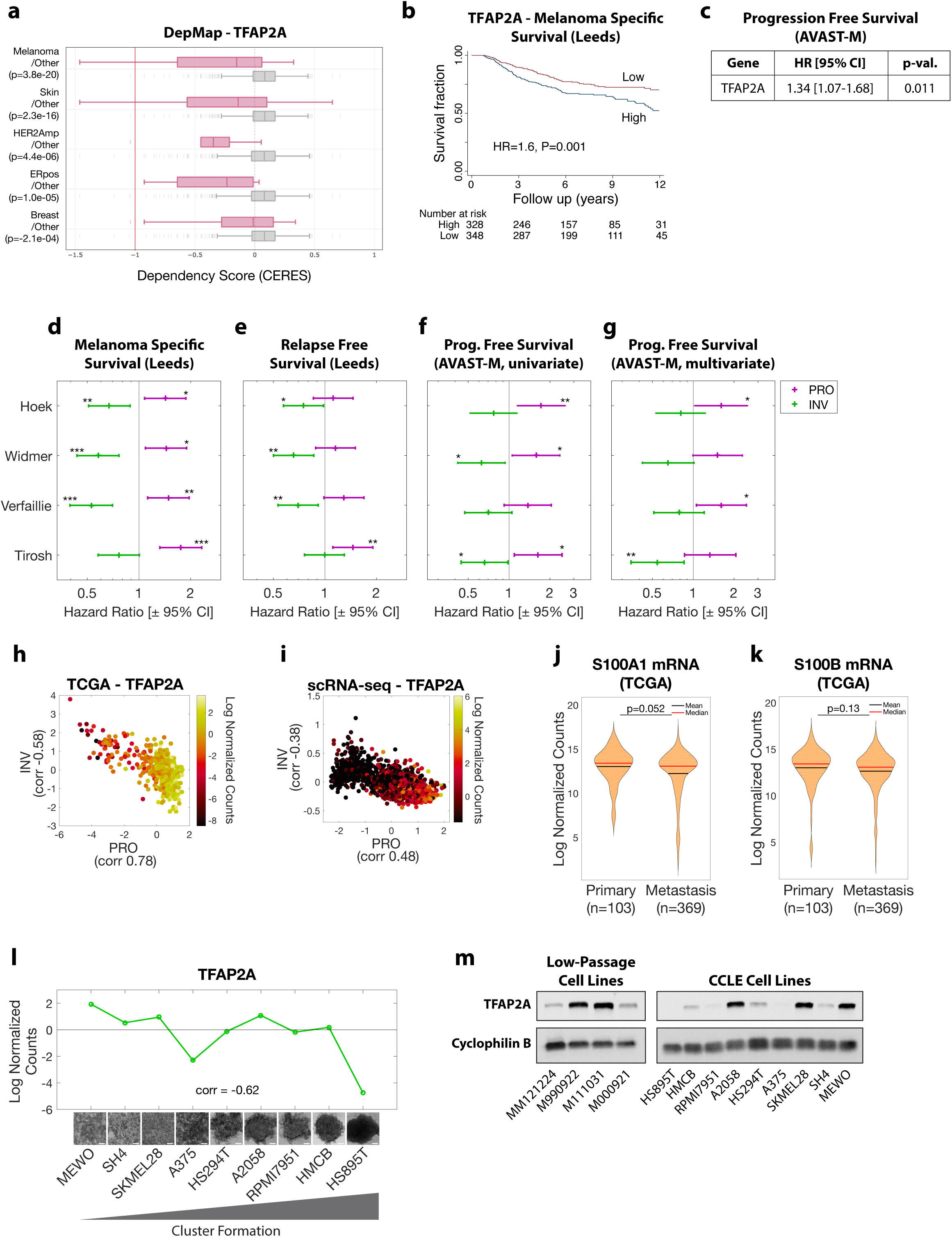
(Related to Figure 5) **a**. Analysis of TFAP2A in the Dependency Map (DepMap, CRISPR (Avana) Public 19Q3) dataset reveals a dependence of melanoma proliferation on TFAP2A. **b-c**. High TFAP2A mRNA expression in primary tumors predicts worse (b) melanoma specific survival in patients in the Leeds Melanoma Cohort (HR [95% CI]: 1.6 [1.2, 2.1], p=0.001 upper vs. lower half by Cox proportional hazards and (c) progression free survival in patients in the AVAST-M Melanoma Cohort (multivariate Cox regression model). **d-g**. Primary tumors with high PRO or low INV expression are associated with worse outcomes in patients in (d-e) the Leeds Melanoma Cohort and (f-g) the AVAST-M Melanoma Cohort. **h**. Human melanoma samples from The Cancer Genome Atlas (TCGA) melanoma (SKCM, n=472) cohort are plotted as PRO and INV scores (Hoek et al., 2006) calculated from RNA-seq and colored according to TFAP2A mRNA expression. Pearson correlation coefficients between TFAP2A and PRO/INV scores are shown on axes. **i**. Individual human melanoma cells are plotted as PRO and INV scores (Hoek et al., 2006) calculated from single-cell RNA-seq and colored according to TFAP2A mRNA expression (re-analyzed from Jerby-Arnon et al. (Jerby-Arnon et al., 2018)). Pearson correlation coefficients between TFAP2A and PRO/INV scores are shown on axes. **j-k**. (j) S100A1 and (k) S100B mRNA expression in The Cancer Genome Atlas (TCGA) melanoma (SKCM) cohort comparing primary tumors and metastases (p-values by Wilcoxon rank sum test with Bonferroni correction). **l**. Human melanoma cell lines ranked by cluster forming ability (left to right: low to high) demonstrate negative correlation between TFAP2A mRNA expression by RNA-seq and clustering (Spearman correlation shown). **m**. Western blot analysis of TFAP2A expression in panels of (left) low-passage human melanoma cell lines and (right) human melanoma cell lines.

**Figure S7.**
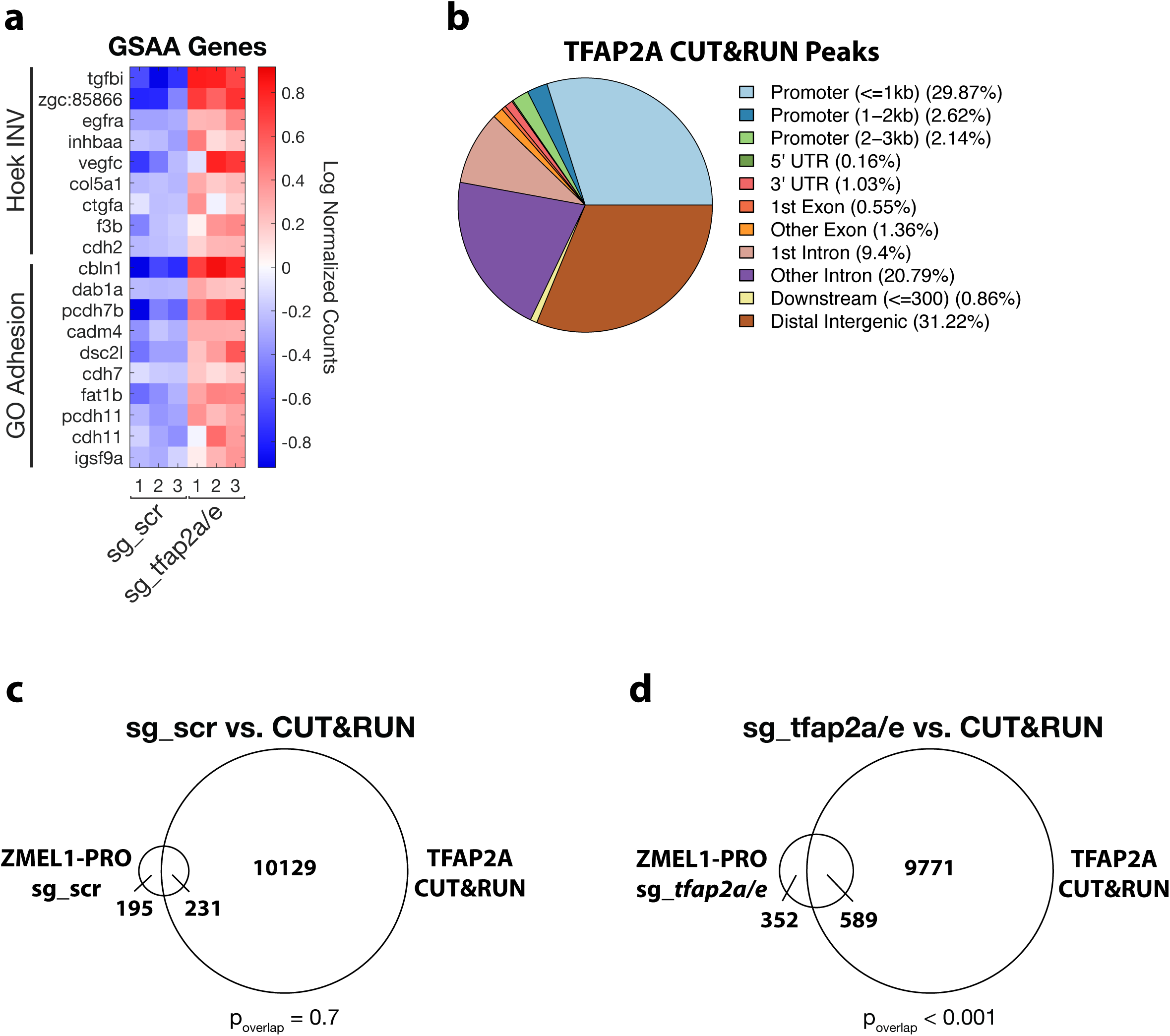
(Related to Figure 5) **a**. Heatmap of top genes in Hoek INV and GO Adhesion gene sets that are differentially expressed between ZMEL1-PRO sg_*tfap2a/e* and sg_scr (log_2_ fold change cutoff ± 0.5, p_adj_ < 0.05). As in Figure 5f, but with zebrafish gene names. **b**. Distribution of TFAP2A CUT&RUN peaks as annotated by ChIPSeeker. **c-d**. Overlap of TFAP2A CUT&RUN peaks with genes upregulated in ZMEL1-PRO following CRISPR/Cas9 with (c) sg_scr (p=0.7 by bootstrapping) and (d) sg_*tfap2a/e* (p<0.001 by bootstrapping).

**Figure S8.**
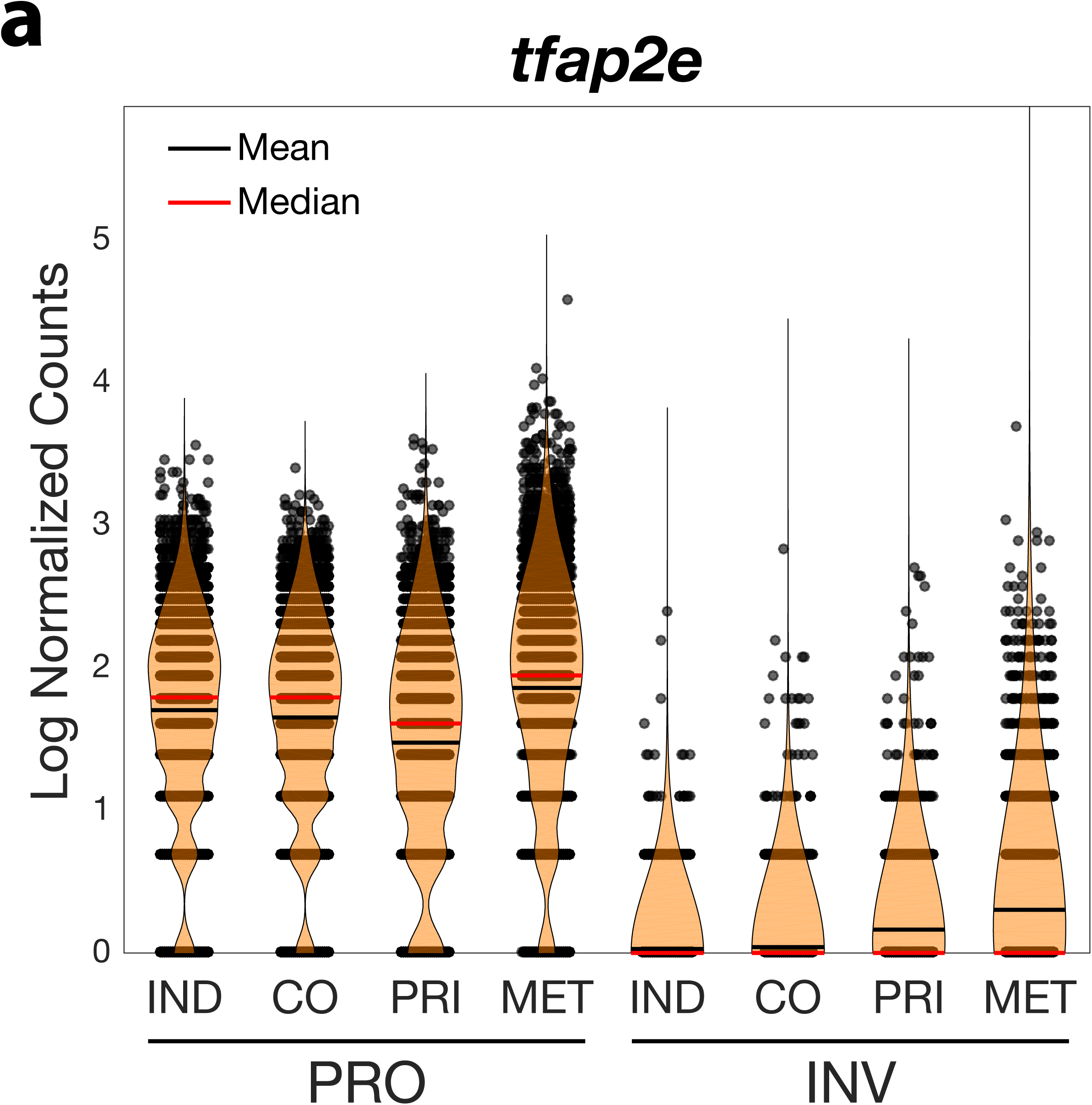
(Related to Figure 6) **a**. Single-cell expression of *tfap2e* in ZMEL1-PRO and -INV cells. Individual culture (IND); co-culture (CO); primary tumors (PRI); metastases (MET).

## Graphical Abstract

Individual melanoma tumors are comprised of proliferative (PRO) and invasive (INV) subpopulations that coexist, each with a set of associated phenotypes. TFAP2 acts as a master regulator of the PRO vs. INV states and clustering, positively regulating melanoma proliferation while negatively regulating both motility/extravasation and clustering. The interaction of these two populations in clusters leads to cooperation in the seeding of metastasis, promoting the formation of heterogenous metastases via collective invasion.

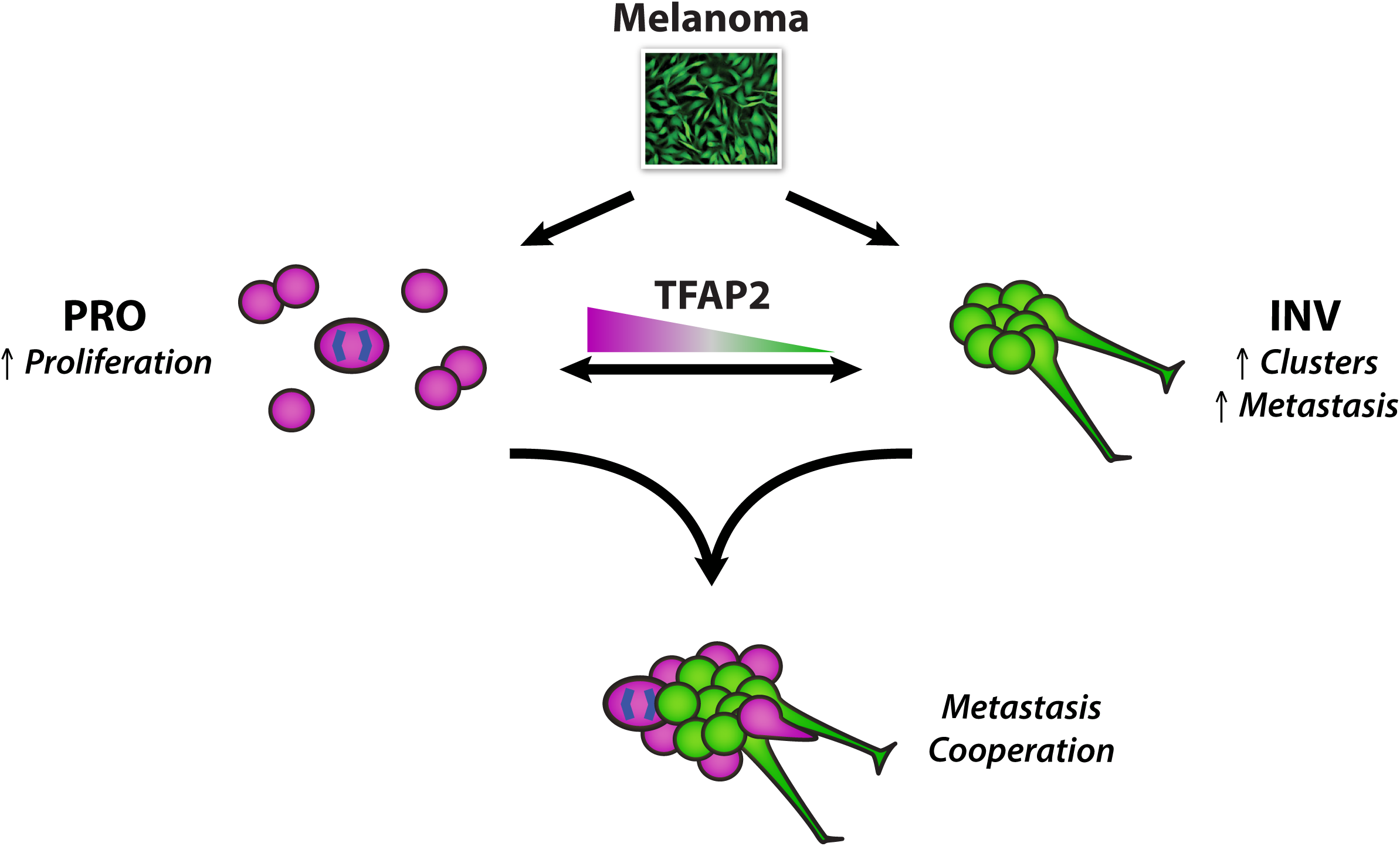

---

**Supplementary Table 1** ZMEL1-INV vs. -PRO RNA-seq results. Filtered differential expression (DE; absolute log_2_ fold change ≥ 1.5, p_adj_ < 0.05); all DE, log normalized counts, and raw counts.

**Supplementary Table 2** ZMEL1 3D vs. 2D culture RNA-seq results for ZMEL1-PRO and ZMEL1-INV. Filtered differential expression (DE; absolute log_2_ fold change ≥ 1.5, p_adj_ < 0.05); all DE, log normalized counts, and raw counts.

**Supplementary Table 3** Annotation of top HOMER motif for 1000bp region (transcription start site ± 500bp) associated with genes upregulated in ZMEL1-PRO. Annotation presented for genes differentially expressed in ZMEL1-PRO and -INV.

**Supplementary Table 4** Annotation of top HOMER motif for 1000bp region (transcription start site ± 500bp) associated with genes differentially expressed in 3D vs. 2D culture for each ZMEL1-PRO and ZMEL1-INV. Annotations presented for top motif for each ZMEL1-PRO and ZMEL1-INV.

**Supplementary Table 5** ZMEL1-PRO sg_*tfap2a/e* vs. sg_scr RNA-seq results. Filtered differential expression (DE; absolute log_2_ fold change ≥ 0.5, p_adj_ < 0.05); all DE, log normalized counts, and raw counts.

**Supplementary Table 6** List of all gene sets used for GSAA, including sources and full gene lists.

**Supplementary Table 7** Results from GSAA analyses. Results table and details for gene sets with false discovery rate (FDR) below 0.05 (up to 6 per analysis).

**Supplementary Table 8** Sequences for sgRNAs used for CRISPR/Cas9 experiments.

**Supplementary Table 9** Full zebrafish to human ortholog mapping table. Orthologs with DIOPT Score < 2 were excluded. A DIOPT Score greater than 6 was considered sufficient for use in GSAA.

**Supplementary Video 1 (Related to Fig. 1h)** Time lapse confocal microscopy of ZMEL1 cells transplanted intravenously in larval zebrafish. Arrowhead indicates group of cells invading from the notochord (NC) and caudal hematopoietic tissue (CHT) into the tail fin mesenchyme (TF).

**Supplementary Video 2 (Related to Fig. 2d)** Time lapse microscopy of ZMEL1 cluster formation in low-bind plates over 3 days.

**Supplementary Video 3 (Related to Fig. 2f)** Time lapse microscopy of cluster formation of 1:1 mixture of ZMEL1-PRO and -INV in low-bind plates over 3 days.

**Supplementary Video 4 (Related to Fig. 3a)** Time lapse confocal microscopy following intravenous transplant of ZMEL1-PRO and -INV. A mixed cluster of both ZMEL1-PRO and -INV populations (11h:00m, arrowhead) extravasated from the caudal hematopoietic tissue (CHT) into the tail fin mesenchyme (TF)—with ZMEL1-INV leading (19h:00m, arrow) and ZMEL1-PRO following (23h:40m, arrow).

## METHODS

### Cloning

Zebrafish-specific expression plasmids were generated by Gateway Cloning (Fisher) into the pDestTol2pA2 backbone (Tol2kit, plasmid #394) (Kwan et al., 2007). nls-mCherry (Tol2kit, plasmid #233) was cloned under the ubiquitin promoter. tdTomato was cloned under the zebrafish *mitfa* promoter.

### Cell culture

The establishment of the ZMEL1 zebrafish melanoma cell line from a tumor in a mitfa-BRAF^V600E^/p53^-/-^ zebrafish was described previously (Heilmann et al., 2015; Kim et al., 2017). ZMEL1 was grown at 28°C in a humidified incubator in DMEM (Gibco #11965) supplemented with 10% FBS (Seradigm), 1X penicillin/streptomycin/glutamine (Gibco #10378016), and 1X GlutaMAX (Gibco #35050061). The ZMEL1-PRO and -INV populations were identified based on phenotyping of multiple concurrent ZMEL1 cultures. ZMEL1 populations were validated by RNA-seq confirming expression of expected transgenes. Human melanoma cell lines were maintained in DMEM (Gibco #11965) supplemented with 10% FBS (Seradigm), 1X penicillin/streptomycin/glutamine (Gibco #10378016), with the exception of HMCB, which was maintained in MEM (Gibco #11095080) supplemented with 10% FBS (Seradigm), 1% Sodium Pyruvate (Gibco #11360070), 1% MEM-Non-Essential Amino Acids (Gibco #11140050), 10mM HEPES (Gibco #15630080), and 1X penicillin/streptomycin (Gibco #15140122). Low-passage human melanoma cell lines were established and cultured as previously described (Raaijmakers et al., 2015). Cells were routinely confirmed to be free from mycoplasma (Lonza Mycoalert). Human cell lines were either purchased directly from ATCC or verified by STR profiling.

### Fluorescently labeled cell lines

The ZMEL1 cell line constitutively expresses EGFP under the *mitfa* promoter (Heilmann et al., 2015). ZMEL1 lines additionally expressing nls-mCherry under the ubiquitin promoter for cell tracking experiments were generated with the Neon Transfection System (Fisher) followed by FACS sorting. ZMEL1 lines expressing tdTomato under the *mitfa* promoter were generated through CRISPR/Cas9 mutation of the constitutive EGFP (see CRISPR/Cas9 below) followed by Neon Transfection and FACS sorting.

### CRISPR/Cas9

The Alt-R CRISPR-Cas9 System (Integrated DNA Technologies) was used for CRISPR-Cas9 experiments following the manufacturer’s protocols for use with the Neon Transfection System (Fisher) and adherent cells. Successful nucleofection was confirmed by visualizing ATTO 550 labeled tracrRNA one day post-nucleofection. Successful loss of full-length protein expression was verified by visualizing loss of EGFP expression (EGFP) or by Western blot (*cdh1, tfap2a, tfap2e*). Control scramble sgRNA (sg_scr) sequence was used from Wang et al. (Wang et al., 2015). sgRNA sequences are listed in Supplementary Table 8.

### Cluster formation assay

ZMEL1 cells were trypsinized, centrifuged at 300g for 3 minutes, and resuspended in standard culture media. A Corning Ultra-Low Attachment Surface 96-well plate (#3474) was seeded with 5×10^4^ cells/well in a final volume of 100uL. Clusters were allowed to form over the course of 2-3 days in a humidified 28°C incubator, or were used for time-lapse microscopy on a Zeiss AxioObserver Z1 equipped with an incubation chamber using a 5x/0.16NA objective. For human melanoma cell lines, a round bottom Corning Ultra-Low Attachment Surface 96-well plate (#7007) was seeded with 5×10^3^ cells/well in a final volume of 100uL. Clusters were allowed to form over the course of 24-48h in a humidified 37°C incubator. Cell lines were ranked according to their relative abilities to form dense three-dimensional clusters.

### Cluster quantification

The average cluster size of each image was quantified using a MATLAB implementation of the characteristic length scale equation from Smeets et al (Smeets et al., 2016). Cluster mixing and spatial sorting were quantified for each cluster using a custom MATLAB segmentation routine. To quantify cluster mixing, background corrected images for EGFP and tdTomato were each segmented using a low and high threshold to improve detection of small and large clusters, respectively. Segmentation from each channel was merged, and each cluster (filtered to have a size corresponding to a size of approximately 4 cells or larger) was classified as red-only, green-only, or red-green mix. Cluster spatial sorting was calculated for individual large (equivalent diameter greater than 45μm) mixed red-green clusters by calculating the weighted average of radial intensity profiles for each channel. The difference between the weighted averages for each channel was calculated and normalized by the radius of the cluster. With this dimensionless metric, small values correspond to well-mixed clusters, whereas larger values correspond to a high degree of spatial segregation.

### Cluster confocal microscopy

ZMEL1 clusters were fixed at room temperature for 45 minutes by adding an equal volume of 4% PFA in PBS to 2-day cultures of clusters (2% final PFA concentration). Fixed clusters were washed once with PBS and counterstained with Hoechst 33342 (Fisher H1399) and transferred to a 96-well glass-bottom plate (Mattek, PBK96G-1.5-5-F) before imaging. Individual clusters were imaged on a Leica TCS SP5-II inverted point-scanning confocal microscope with a or 40x/1.10NA objective. 3D reconstruction was performed using Volocity (PerkinElmer, v6.3). Individual slices were visualized with ImageJ.

### Cell tracking

Time-lapse microscopy was performed on a Zeiss AxioObserver Z1 equipped with an incubation chamber. A 96-well plate (Corning 353072) was seeded with 1.2×10^4^ ZMEL1 cells (PRO or INV) admixed such that 1/6 of the population stably expressed nls-mCherry under the Ubi promoter, and allowed to adhere overnight. Cells were imaged every 5 minutes for 24 hours. Centroids of nuclei were identified, and tracks generated using a MATLAB implementation of the IDL tracking methods developed by John Crocker, David Grier, and Eric Weeks (physics.georgetown.edu/matlab/). For each imaging location, growth was calculated based on the number of nuclei present at each time point, assuming equal numbers of cells move in and out of the field of view over time. For each track, mean squared displacement (MSD) was calculated as previously described (Gorelik and Gautreau, 2014) over a range of lag times (5≤τ<100 min). The log-log plot of MSD vs. τ provides information about both the diffusion coefficient (intercept) and persistence (slope, α) of cells. For a cell moving randomly, α=1; and for a cell moving along a straight line, α=2 (Gorelik and Gautreau, 2014). N=4 independent replicates were performed, each consisting of 6 technical replicates per cell type. Growth rates were quantified with a linear mixed-effects model using the fitlme function in MATLAB with the model, ‘log(cell_number) ∼ time + cell_type:time + (1. replicate)’. Motility (α) was quantified as the slope of the linear model, ‘log(MSD) ∼ 1 + cell_type*log(τ)’. Growth plots represent the smoothed (moving window average of 5 time points [= 25 min]) average cell number ± SE normalized to the cell number at time zero.

### Boyden chamber migration

Cell migration of PRO/PRO, INV/INV, and PRO/INV mixtures was quantified using a Boyden chamber (transwell) assay. A 1:1 mixture of ZMEL1 cells labeled with EGFP and tdTomato (5×10^4^ cells total per well) were added to each transwell insert (Corning, 353492, 3.0um pore) in a 24-well plate (Corning, 353504) in 500uL of complete media. The lower chamber was filled with 500uL of complete media and cells were allowed to migrate for 2 days. Cells were fixed for 15 minutes at room temperature with 4% PFA in PBS, washed once with PBS, and non-migrated cells removed with cotton-tipped swabs. Migrated cells were stained with Hoechst 33342 (Fisher H1399) and ≥9 fields/well imaged on a Zeiss AxioObserver Z1 with a 10x/0.45NA objective. Nuclei were segmented using intensity thresholding of background-corrected Hoechst staining followed by an intensity-based watershed step to separate adjacent objects. Cell identity was established based on fluorophore expression within the mask defining each nucleus.

### Conditioned media

Conditioned media was collected from confluent 10cm dishes of ZMEL1-PRO and -INV cultures following 2-3 days of growth and filtered through a 0.45μm syringe filter (Fisher #09-720-005) to remove viable cells. Filtered conditioned media or fresh complete media was mixed 1:1 with fresh complete media and used for subsequent assays. For proliferation assays, ZMEL1-PRO or -INV cells (1.4×10^4^ cells/well) were plated in white wall 96-well plates (Corning #3610) in a 1:1 mixture of fresh complete media with either ZMEL1-PRO or -INV conditioned media or fresh complete media. Relative proliferation was measured by CellTiter-Glo® Luminescent Cell Viability Assay (Promega) according to manufacturer’s protocol two days after plating. For Boyden Chamber assays, ZMEL1-PRO or -INV cells (5×10^4^ cells total per well) were added to each transwell insert (Corning, 353492, 3.0um pore) in a 24-well plate (Corning, 353504) in 500uL of fresh complete media. The lower chamber was filled with either ZMEL1-PRO or -INV conditioned media or fresh complete media and cells were allowed to migrate for 2 days. Non-migrated cells were removed with cotton-tipped swabs. Migrated cells were stained with Hoechst 33342 (Fisher H1399) and 26 fields/well imaged on a Zeiss AxioObserver Z1 with a 10x/0.45NA objective. Centroids of nuclei were identified and counted using a MATLAB implementation of the IDL tracking methods developed by John Crocker, David Grier, and Eric Weeks (physics.georgetown.edu/matlab/).

### Zebrafish husbandry

Zebrafish were housed in a dedicated facility maintained at 28.5°C with a light/dark cycle (14 hours on, 10 hours off). All anesthesia was performed using Tricaine-S (MS-222, Syndel USA, Ferndale, WA) with a 4g/L, pH 7.0 stock. All procedures adhered to Memorial Sloan Kettering Cancer Center IACUC protocol number 12-05-008.

### Larval transplantation

Transplantation of ZMEL1 cells into 2dpf *casper* zebrafish larvae was performed as previously described (Heilmann et al., 2015; Kim et al., 2017). Briefly, ZMEL1 cells were prepared by trypsinization, centrifuged at 300g for 3 minutes, and resuspended at a concentration of either 2.5×10^7^ or 5.0×10^7^ cells/mL in 9:1 DPBS:H_2_O (Gibco 14190-144). Cells were injected into the Duct of Cuvier of 2dpf *casper* or *casper* FLK-RFP (labeling the vasculature with RFP) fish using a Picoliter Microinjector (Warner Instruments, PLI-100A) with a glass capillary needle (Sutter, Q100-50-10) made on a laser-based needle puller (Sutter, P-2000). For mixing studies, ZMEL1-PRO and ZMEL1-INV differentially labeled with EGFP or tdTomato were mixed at a 1:1 ratio prior to injection. Fish with successful transplants based on the presence of circulating cells and/or cells arrested in the caudal vasculature were either used for time-lapse confocal microscopy (see “Zebrafish confocal time-lapse imaging”) or individually housed and followed by daily imaging on a Zeiss AxioZoom V16 fluorescence microscope.

### Adult transplantation

Transplantation of ZMEL1 cells into adult *casper* zebrafish was performed as previously described (Heilmann et al., 2015; Kim et al., 2017). Briefly, adult zebrafish were irradiated on two sequential days with 15 Gy on a cesium irradiator (Shepherd) and were transplanted 3-4 days later. On the day of transplant, ZMEL1 cells were prepared by trypsinization, washed once with DPBS, and resuspended at a concentration of 1.67×10^8^ cells/mL in DPBS (Gibco 14190-144). Cells were injected subcutaneously, caudal to the cloaca, on the ventral side of zebrafish anesthetized with Tricaine-S. Fish were imaged on days 1, 3, and 7 post-transplant on a Zeiss AxioZoom V16 fluorescence microscope. For 1:1 mixing studies, fish were injected with an equivalent final concentration of either ZMEL1-PRO, ZMEL1-INV, or a 1:1 mixture of the two populations differentially labeled with EGFP or tdTomato. A total of N=5 independent mixing experiments were performed, N=3 for INV-EGFP/PRO-tdTomato and N=2 for INV-tdTomato/PRO-EGFP, each with at least 12 fish per group (n=231 fish total). For variable ratio mixing studies, fish were injected with an equivalent final concentration of tdTomato-labeled ZMEL1-PRO mixed at a 1:4, 4:1, or 9:1 ratio with EGFP-labeled ZMEL1-PRO or ZMEL1-INV. A total of N=3 independent variable ratio mixing experiments were performed, each with at least 16 fish per group (n=334 fish total).

### Zebrafish imaging and image quantification

#### Whole-fish larval imaging

Larval zebrafish transplanted as described above were anesthetized with Tricaine-S and imaged on a bed of 2% agarose (KSE Scientific, BMK-A1705) in E3. Images were manually scored for cells that had invaded the tail fin parenchyma at experiment endpoint.

#### Adult imaging

Adult zebrafish were imaged as previously described. (Heilmann et al., 2015) Briefly, on days 1, 3, and 7 post-transplant fish were anesthetized with Tricaine-S and imaged on a bed of 2% agarose using a monocolor camera for fluorescence and brightfield, and color camera for observing pigmentation. Fluorescence images were manually scored for the following pre-specified binary outcomes:

- Distant metastases: tumor not adjacent or contiguous with primary tumor
- Caudal metastases: distant metastasis caudal to anal fin
- Dorsal metastases: distant metastasis on dorsum of fish, near insertion of dorsal fin

For each binary outcome, the population composition was also scored (PRO, INV, or both). Primary tumor growth was also quantified over time for each fish using a previously described custom MATLAB pipeline (Heilmann et al., 2015). An adapted version of this pipeline with an adaptive threshold segmentation was utilized to allow easier visualization of representative images.

### Zebrafish larval confocal time-lapse imaging

Larval zebrafish were transplanted as described above. Fish with successful intravenous transplants were anesthetized with Tricaine-S and embedded in 1% low-melt agarose (Sigma A9045) in E3 containing 0.28ug/mL Tricaine-S in a glass-bottom square-well 96-well plate (Arrayit 96-well Microplate SuperClean, Cat M96FC, Lot 150901). Wells were filled with E3 containing 0.28ug/mL Tricaine-S and the plate sealed with a PCR microseal (BioRad Microseal ‘B’ Film, #MSB1001). Up to 45 larval zebrafish per experiment were imaged in parallel on a GE IN Cell Analyzer 6000 every 15-20 minutes for 24-30h. A single z-stack was acquired for each fish using a 10X/0.45NA objective and 8-10um z-steps. Because the ventral edge of the caudal vein exists as a single plane at this developmental stage (2-3dpf) (Isogai et al., 2001), maximum intensity projections were generated for each fish and manually scored for the presence of ZMEL1 cells that extravasate ventrally from the caudal vein and invade into the tail fin. Tracking of collectively migrating clusters in ZMEL1-INV cells labeled with nls-mCherry was performed with the MTrackJ (Meijering et al., 2012) plugin in ImageJ.

### Western blot

Cell lysates were collected by sonication in RIPA buffer (Thermo #89901) with 1X Halt Protease and Phosphatase Inhibitor Cocktail (Thermo #78441) followed by centrifugation (14,000rpm for 10min at 4°C) and collection of the supernatant. Protein concentration was quantified by Bradford (Sigma B6916-500mL) according to manufacturer’s protocol. Samples were mixed with 6X reducing loading buffer (Boston BioProducts #BP-111R) and denatured at 95°C for 10 minutes. Samples were run on a Mini PROTEAN TGX gel (BioRad) and transferred using Turbo Mini Nitrocellulose Transfer Pack (Bio-Rad, catalog #1704158). Membranes were blocked with 5% nonfat dry milk in TBST (1X TBS + 0.1% Tween 20) for 1 hour before incubation with primary antibody in PBS overnight at 4°C. Membranes were washed with TBST and incubated with secondary antibody in 5% nonfat dry milk for 1 hour at room temperature. Membranes were washed with TBST and developed with ECL (Amersham, RPN2109) using an Amersham Imager 600 (GE) or chemiluminescence film. Antibodies: anti-hs_CDH1 (BD #610181, lot 8082613), anti-dr_Tfap2a (LifeSpan Biosciences, #LS-C87212, log 113877), anti-dr_Tfap2e (Fisher, #PA5-72631, lot UA2709682A), anti-hs_TFAP2A (Cell Signaling, #3215, clone C83E10, lot 2), anti-hs_cyclophilin B (Fisher #PA1-027A, lot SD248938).

### Immunofluorescence

Cells were allowed to adhere for 2 days to glass chamber slides (PEZGS0816) coated with poly-D-lysine (Sigma #P-0899). Slides were fixed for 15 minutes with 4% PFA in PBS, washed 3 times with PBS, and permeabilized with 0.1% Triton X-100 (Fisher #BP151-100) in PBS for 15 min. Slides were blocked with 5% donkey serum (Sigma #S30-M), 1% BSA (Fisher #BP1600-100), and 0.1% Triton X-100 in PBS for 1 hour, followed by incubation in primary antibody overnight at 4°C. Slides were washed 3 times with PBS and incubated in secondary antibody (anti-mouse [Cell Signaling, #4408S]; anti-rabbit [Cell Signaling, #8889S]) for 1.5 hours, followed by 3 washes with PBS and counterstain with (1:10,000) Hoechst 33342 in PBS for 45 minutes. Slides were washed with PBS and mounted in ProLong Glass Antifade Mountant (Fisher #P36984). All incubations were carried out at room temperature unless otherwise noted. Antibodies: rabbit anti-hs_TFAP2A (Cell Signaling, #3215, clone C83E10, lot 2), mouse anti-hs_H3 (Cell Signaling, #14269, clone 1B1B2, lot 6). Slides were imaged in a minimum of 9 fields on a Zeiss AxioObserver Z1 using a 20x/0.80NA objective. Nuclei were segmented using intensity thresholding of background-corrected Hoechst staining followed by an intensity-based watershed step to separate adjacent objects. TFAP2A expression was quantified for each nucleus as the ratio of TFAP2A to Histone H3 staining intensity.

### RNA-seq

#### Samples

For 2D culture, three replicate cultures at 70-80% confluence for each ZMEL1-PRO and -INV were utilized. For 3D culture, three replicate cultures for each ZMEL1-PRO and -INV grown in Corning Ultra-Low Attachment Surface 6-well plates (Corning #3471) for 48 hours were utilized. One individual replicate of RNA-seq of ZMEL1-PRO grown in 3D culture was excluded due to low RIN score, poor SeQC metrics and poor clustering with other replicate samples. For *tfap2a/e* CRISPR, three independent batches of ZMEL1-PRO cells nucleofected with either sg_scr or sg_*tfap2a/e* and grown in 2D conditions were utilized at 8-16 days post nucleofection.

#### Sequencing and analysis

Total RNA was extracted with the RNeasy Plus Mini kit with QiaShredder (Qiagen). Purified RNA was delivered to GENEWIZ (South Plainfield, NJ) for mRNA preparation with the TruSeq RNA V2 kit (Illumina) and 100bp (2D) or 150bp (3D and CRISPR) paired-end sequencing on the Illumina HiSeq2500. After quality control with FASTQC (Babraham Bioinformatics) and trimming with TRIMMOMATIC (Bolger et al., 2014) when necessary, reads were aligned to GRCz10 (Ensembl version 81) using STAR (Dobin et al., 2013), with quality control via SeQC (DeLuca et al., 2012). Differential expression was calculated with DESeq2 (Love et al., 2014) using the output of the --quantMode GeneCounts feature of STAR. The rlog function was used to generate log_2_ transformed normalized counts. Pathway and Gene Ontology (GO) analysis were performed with GSAA using the following parameters: gsametric Weighted_KS, demetric Signal2Noise, permute gene_set, rnd_type no_balance, scoring_scheme weighted, norm MeanDiv (Xiong et al., 2012; Xiong et al., 2014). A full list of gene sets used for GSAA can be found in Supplementary Table 6. Ortholog mapping between zebrafish and human was performed with DIOPT (Hu et al., 2011) (Supplementary Table 9). Only orthologs with a DIOPT score greater than 6 were used for GSAA and heatmap generation. In cases of more than one zebrafish ortholog of a given human gene, the zebrafish gene with the highest average expression was selected. De-novo motif analysis was performed with the HOMER (Heinz et al., 2010) function findMotifs.pl, using the zebrafish genome (GRCz10) and searching for motifs of lengths 8, 10, and 16 within ± 500bp of the TSS of differentially expressed genes. Motifs were annotated using JASPAR (Khan et al., 2017). A Cell-Cell Adhesion gene set was defined from the Core Enrichment genes from comparing ZMEL1-INV vs. -PRO to the gene set, *GO Cell Cell Adhesion Via Plasma Membrane Adhesion Molecules*.

### TFAP2A CUT&RUN

#### Sample preparation

Anti-TFAP2A Cleavage Under Targets and Release Using Nuclease (CUT&RUN) sequencing was performed in wild-type and *TFAP2A*;*TFAP2C* double-mutant SKMEL28 cell lines as described (Skene and Henikoff, 2017) with minor modifications. Cells in log-phase culture (approximately 80% confluent) were harvested by cell scraping (Corning), centrifuged at 600g (Eppendorf, centrifuge 5424) and washed twice in calcium-free wash-buffer (20 mM HEPES, pH7.5, 150 mM NaCl, 0.5 mM spermidine and protease inhibitor cocktail, cOmplete Mini, EDTA-free Roche). Pre-activated Concanavalin A-coated magnetic beads (Bangs Laboratories, Inc) were added to cell suspensions (2×10^5^ cells) and incubated at 4°C for 15 mins. Antibody buffer (wash-buffer with 2mM EDTA and 0.03% digitonin) containing anti-TFAP2A (abcam, ab108311) or Rabbit IgG (Millipore, 12-370) was added and cells were incubated overnight at 4°C. The next day, cells were washed in dig-wash buffer (wash buffer containing 0.025% digitonin) and pA-MNase was added at a concentration of 500 μg/ mL (pA-MNase generously received from Dr. Steve Henikoff). The pA-MNase reactions were quenched with 2X Stop buffer (340mM NaCl, 20mM EDTA, 4mM EGTA, 0.05% Digitonin, 100 μg/ mL RNAse A, 50 μg/ mL Glycogen and 2 pg/ mL sonicated yeast spike-in control). Released DNA fragments were Phosphatase K (1μL/mL, Thermo Fisher Scientific) treated for 1 hr at 50°C and purified by phenol/chloroform-extracted and ethanol-precipitated. Fragment sizes analysed using an 2100 Bioanalyzer (Agilent). All CUT&RUN experiments were performed in duplicate.

#### Library preparation and data analysis

CUT&RUN libraries were prepared using the KAPA Hyper Prep Kit (Roche). Quality control post-library amplification was conducted using the 2100 Bioanalyzer (Agilent) for fragment analysis. Libraries were pooled to equimolar concentrations and sequenced with paired-end 100 bp reads on an Illumina HiSeq X platform. Paired-end FastQ files were processed through FastQC (Babraham Bioinformatics) for quality control. Reads were trimmed using Trim Galore Version 0.6.3 (Developed by Felix Krueger at the Babraham Institute) and Bowtie2 version 2.1.0 (Langmead and Salzberg, 2012) was used to map the reads against the hg19 genome assembly. The mapping parameters and peak calling analysis was performed as previously described (Meers et al., 2019). Called peaks were annotated with ChIPseeker v1.18.0 (Yu et al., 2015), with Distal Intergenic peaks excluded from downstream analysis. P-values for overlap with differential expression gene sets were calculated by comparing against overlap with randomly selected gene sets (n=10,000 iterations).

### Single-cell RNA-seq (scRNA-seq)

#### Sample preparation

*In vitro* samples were cultured under standard conditions with either ZMEL1-PRO (tdTomato) and -INV (EGFP) separately (individual culture) or mixed together in a 1:1 ratio for 11 days (co-culture). Cells were trypsinized, resuspended in DMEM supplemented with 2% FBS, and flow sorted using a SY3200 (Sony) for DAPI-negative pure ZMEL1-PRO and -INV populations prior to droplet-based scRNA-seq. For *in vivo* samples, adult *casper* zebrafish were transplanted with a 1:1 mixture of ZMEL1-PRO (tdTomato) and -INV (EGFP) cells as described above (Adult transplantation). Tumors were allowed to grow for 6 days (primary tumors) or 13 days (metastases). At experimental timepoint, tumors were surgically excised and minced with a fresh scalpel (primary tumors from n=6 fish; metastases from n=4 fish). Each sample was placed in a 15mL tube containing 3mL of 0.9X DPBS with 0.16 mg/mL of Liberase TL (Sigma #5401020001), incubated at room temperature for 15 minutes followed by trituration using a wide-bore P1000 (Fisher #2069G), and incubated for an additional 15 minutes. 500μL FBS was added and each sample was triturated again and passed through a 70μm cell strainer (Corning #352350). Samples were centrifuged 500g for 5 minutes, resuspended in DMEM supplemented with 2% FBS, and flow sorted using a SY3200 (Sony) for DAPI-negative pure ZMEL1-PRO and -INV populations prior to droplet-based scRNA-seq.

#### Droplet-based scRNA-seq

For droplet-based scRNA-seq, experiments were performed using the 10X Genomics Chromium platform, with the Chromium Single Cell 3′ Library & Gel Bead Kit v3.1 (1000128) and Chromium Single Cell 3′ Chip G (1000127). ∼8,000 cells per condition were centrifuged 300g for 5 minutes and resuspended in DMEM supplemented with 10% FBS and loaded to each channel for GEM generation and barcoded single cell libraries were generated according to manufacturer’s instructions. Libraries were diluted to 2nM and 75bp paired end sequencing was performed using the Illumina NextSeq 500. Between 150-200 million paired reads were generated for each library.

#### scRNA-seq processing and analysis

Raw sequencing data was processed using the CellRanger 3.1.0 pipeline developed by 10X Genomics. First, a custom zebrafish genome was generated based on GRCz10 with the addition of exogenous transgenes EGFP, tdTomato, and human BRAF-V600E using the command *cellranger mkref*. Next, the command *cellranger count* was utilized to perform alignment, filtering, barcode counting and UMI counting of all the samples. The Seurat R package (Version 3.1.4) was used for quality control, normalization and dimensionality reduction. Low quality cells with a) features less than 200 or greater than 5000, b) total counts less than 5000, or c) mitochondrial content greater than 15%, were discarded from the analysis. After filtering, normalization was performed using Seurat’s SCTransform procedure with default parameters to perform a regularized negative binomial regression based on the 3,000 most variable genes. Uniform Manifold Approximation and Projection (UMAP) (McInnes et al., 2018) dimensionality reduction was performed using default parameters. The log transformed normalized count data was extracted from Seurat and used for downstream analysis in MATLAB. Plots of PRO vs. INV scores were generated as described in Tirosh et al. (Tirosh et al., 2016) using PRO and INV gene sets defined from the top 250 most differentially expressed genes in each population by ZMEL1 bulk RNA-seq. Briefly, log-transformed data was mean-centered. For each gene set a control gene set was defined to control for variations in sequencing depth and library complexity by randomly selecting 100 genes from the same expression bin (n=25 bins), such that a 50 gene set would have a control gene set of 5,000 genes. The score for each sample was defined as the mean expression of the gene set minus the mean expression of the respective control gene set. A binary classifier for PRO versus INV state was defined by logistic regression on *in vitro* individual culture samples and used to classify cells from all conditions.

### Analysis of publicly available RNA-seq data

#### Cancer Cell Line Encyclopedia (CCLE)

RNA-seq expression data (RSEM genes TPM, version 20180929) was downloaded from the Broad CCLE (https://portals.broadinstitute.org/ccle). Plots of PRO vs. INV scores and their correlations with TFAP2A mRNA expression were generated as for ZMEL1 scRNA-seq data above using log-transformed [log_2_(TPM+1)] and mean-centered data with PRO and INV gene sets from Hoek et al. (Hoek et al., 2006).

#### The Cancer Genome Atlas (TCGA)

Skin Cutaneous Melanoma (SKCM) mRNA expression (v2 RSEM genes normalized, version 2016_01_28, 472 samples from 469 patients) was downloaded from the Broad GDAC Firehose (http://firebrowse.org/). PRO and INV gene expression scores and their correlations with TFAP2A mRNA expression were calculated and plotted as described for CCLE data above. Log-transformed normalized expression of TFAP2A and the pan-melanoma markers (Gaynor et al., 1981; Xiong et al., 2019) S100A1 and S100B were compared between samples from primary tumors and metastatic sites (Wilcoxon rank sum test with Bonferroni correction).

#### Human melanoma single-cell RNA-seq

Processed data for human melanoma single-cell RNA-seq (Tirosh et al., 2016) was downloaded from NCBI GEO (GSE72056). PRO and INV gene expression scores and their correlations with TFAP2A mRNA expression were calculated and plotted as described for CCLE data above.

### DepMap

TFAP2A was queried through the Broad Institute Dependency Map (DepMap) portal (https://depmap.org/portal/) using the CRISPR (Avana) Public 19Q3 dataset.

### External validation using the AVAST-M melanoma cohort

Bulk RNA-seq data from 194 primary melanoma patients was extracted from the phase III adjuvant AVAST-M melanoma cohort (Corrie et al., 2018; Garg et al., 2020). Variance stabilizing transformation (VST) was applied to the raw counts using the *varianceStabilizingTransformation* function from the package DESeq2 (Love et al., 2014) (v1.22.2).

#### TFAP2A survival analysis

VST normalized expression data was used as a continuous variable in a multivariate Cox regression model, using the coxph function of the survival package (Therneau, 2020; Therneau and Grambsch, 2000) (v2.42-3) in R (v3.5.1). Progression-free survival was calculated as the time from diagnosis to the last follow-up or death/progression to metastatic disease, whichever occurred first. The following clinical covariates were considered in the multivariate Cox regression model; age at diagnosis, gender, stage (AJCC 7^th^ edition), ECOG (Eastern Cooperative Oncology Group Performance Status), NClass (regional lymph involvement) and treatment (bevacizumab or placebo) (Garg et al., 2020).

#### PRO/INV survival analysis

VST expression data corresponding to the genes listed in the PRO and INV gene sets were extracted (Hoek et al., 2006; Tirosh et al., 2016; Verfaillie et al., 2015; Widmer et al., 2012). For each sample, the expression values of these genes were standardized to have zero mean and unit standard deviation. The mean of these standardized expression values was computed to obtain a vector score corresponding to the PRO and INV expression score vectors. These PRO and INV expression score vectors were divided into “high” and “low” expression groups using the median cut-off. Cox regression models were then fitted by means of the coxph function of the survival package (Therneau, 2020; Therneau and Grambsch, 2000) (v2.42-3) in R (v3.5.1). The hazard ratio (HR) (95% CI) and p-values corresponding to the “high” expression score group were reported in both univariate and multivariate analyses.

### External validation using the Leeds Melanoma Cohort

Primary tumor expression of TFAP2A as well as PRO and INV signatures were tested for association with melanoma specific survival and relapse-free survival by Cox proportional hazards in a large population-based cohort (n=703, accession number EGAS00001002922) (Nsengimana et al., 2018). Signature scores were created by averaging z-transformed gene expressions and dichotomized by median split.

### Statistical analysis

Statistical analysis and figure generation were performed in MATLAB (Mathworks, R2016a). RNA-seq analysis was performed in R (R Foundation for Statistical Computing, 3.4.0). Image processing and analysis was performed using MATLAB, Zen (Zeiss), ImageJ (NIH), and Volocity (PerkinElmer, v6.3). The Leeds Melanoma Cohort was analyzed in STATA v14 (StataCorp, Texas, USA). Unless otherwise noted, bar plots represent mean ± standard deviation (SD) of independent experiments, and dots represent means of independent experiments. Abbreviations for p-values are as follows: * p<0.05, ** p<0.01, *** p<0.001.

### Data and reagent availability

All RNA-seq data generated in this study are available via the NCBI GEO repository (GSE151679), with bulk RNA-seq counts and differential expression tables in the Supporting Information. TFAP2A CUT&RUN data are available via the NCBI GEO repository (GSE153020). Cell lines generated in this work are available upon request. Raw data are available upon request.

### Code availability

A MATLAB-based image analysis pipeline for quantifying melanoma in zebrafish was previously published (Heilmann et al., 2015). Additional scripts are available upon request.

